# Fluorescence-Detected Conformational Changes in Duplex DNA in Open Complex Formation by *E. coli* RNA Polymerase: Upstream Wrapping and Downstream Bending Precede Clamp Opening and Insertion of the Downstream Duplex

**DOI:** 10.1101/2020.02.04.932780

**Authors:** Raashi Sreenivasan, Irina A. Shkel, Munish Chhabra, Amanda Drennan, Sara Heitkamp, Hao-Che Wang, Malavika A. Sridevi, Dylan Plaskon, Christina McNerney, Katelyn Callies, Clare K. Cimperman, M. Thomas Record

## Abstract

FRET (fluorescence energy transfer) between far-upstream (−100) and downstream (+14) cyanine dyes showed extensive bending/wrapping of λP_R_ promoter DNA on *E. coli* RNA polymerase (RNAP) in closed and open complexes (CC, OC). Here we determine the kinetics and mechanism of DNA bending/wrapping by FRET and of formation of RNAP contacts with −100 and +14 DNA by single-dye fluorescence enhancements (PIFE). FRET/PIFE kinetics exhibit two phases: rapidly-reversible steps forming a CC ensemble ({CC}c of four intermediates (initial (RP_C_), early (I_1E_), mid-(I_1M_), late (I_1L_)), followed by conversion of {CC} to OC via I_1L_. FRET and PIFE are first observed for I_1E_, not RP_c_. FRET/PIFE together reveal large-scale bending/wrapping of upstream and downstream DNA as RP_C_ advances to I_1E_, reducing −100/+14 distance to ∼75Å and making RNAP-DNA contacts at −100 and +14. We propose that far-upstream DNA wraps on the upper β’-clamp while downstream DNA contacts the top of the β-pincer in I_1E_. Converting I_1E_ to I_1M_ (~1s time-scale) *reduces* FRET efficiency with little change in −100/+14PIFE, interpreted as clamp-opening that moves far-upstream DNA (on β’) away from downstream DNA (on β) to increase the −100/+14 distance by ~14Å. FRET increases greatly in converting I_1M_ to I_1L_, indicating bending of downstream duplex DNA into the clamp and clamp-closing to reduce the −100/+14 distance by ~21Å. In the subsequent rate-determining DNA-opening step, in which the clamp may also open, I_1L_ converts to the initial unstable OC (I_2_). Implications for facilitation of CC-to-OC isomerization by upstream DNA and upstream-binding, DNA-bending transcription activators are discussed.

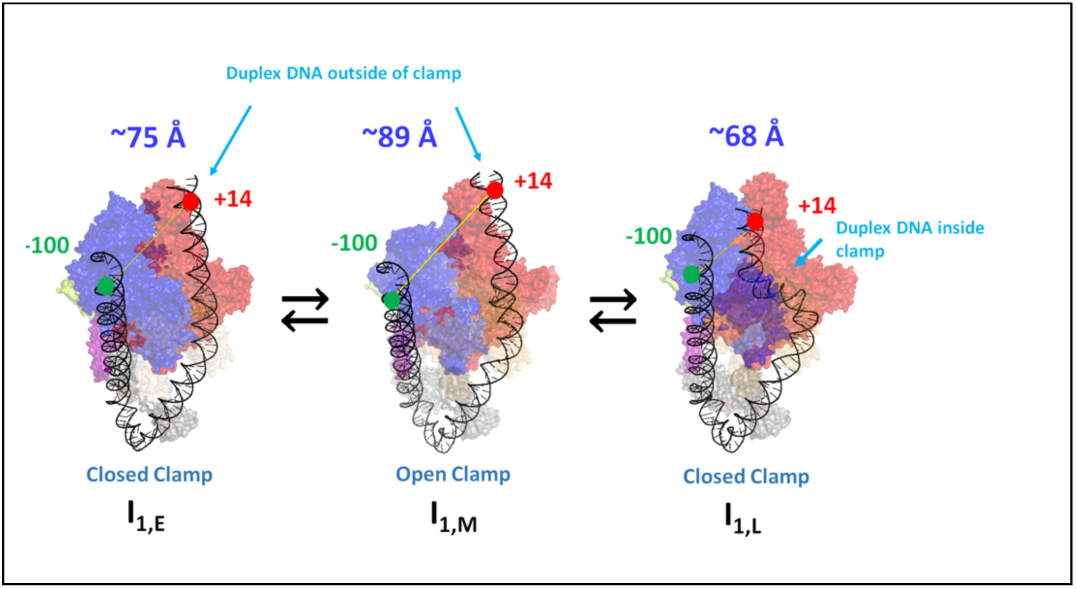

## Introduction

The rate of open complex (OC) formation by *E. coli* RNA polymerase σ^70^ holoenzyme (RNAP; R) at promoter DNA (P) is an important determinant of the rate of transcription initiation.^*1, 2*^ Hence the kinetics and equilibria of the steps of OC formation are highly regulated by promoter sequence, transcription factors, ligands, and solution variables. Monitored by abortive initiation and filter binding assays that detect only long-lived OC, the kinetics of OC formation usually are well-described by the two-step minimal Mechanism 1.^*1-7*^

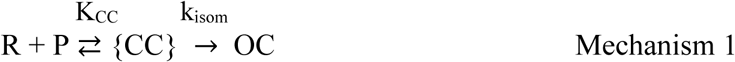

In Mechanism 1, K_CC_ is the composite equilibrium constant for reversible formation of an ensemble of early and advanced closed complexes (called the CC ensemble, symbolized as {CC}) from promoter DNA.^*4*^ Evidence for three likely members of {CC} is provided by the different downstream boundaries of chemical and/or enzymatic CC footprints obtained at different promoters. These CC include the initial specific complex (RP_C_; downstream footprint boundary at approximately −5)^*8-14*^ and two more-advanced CC (designated I_1E_ and I_1L_ for “early” and “late”) with downstream footprint boundaries at approximately +2/+7^*15*^ and at +20.^*8, 9, 11, 14*^ (All positions are numbered relative to the +1 start site.) The isomerization rate constant k_isom_ is the composite rate constant for conversion of {CC} to OC via I_1L_, the most advanced member of {CC}, and the subsequent DNA opening step.^*4*^

This interpretation of the parameters of Mechanism 1 differs subtly but significantly from that used originally. Most previous discussions of effects of changes in promoter sequence or length, transcription factors and solution variables on K_CC_ and k_isom_ have not considered changes in the composition of the equilibrated {CC} ensemble with these variables. In general, any shift in the proportions of the different members of the equilibrated {CC} ensemble will change both K_CC_ and k_isom_ (see ref. 4 and SI Eqs. S19-22). The large effects of upstream truncation of promoter DNA on k_isom_^*16, 17*^ are well-explained in this way (see ref. 4 and Discussion).

The initial closed complex RP_C_ is stabilized by some combination of specific interactions of the flexibly-tethered αCTDs with the promoter UP element (a 20 bp region containing A and T tracts immediately upstream of the −35 region)^*18*^ and of regions 2 and 4 of the σ^70^ subunit with the promoter −10 and −35 elements.^*19*^ After RP_C_ formation, the downstream duplex must bend by at least 90° into the open clamp in order for 13-14 bp, including the −10 and discriminator regions and the transcription start site, to be opened (melted) by RNAP using binding free energy.^*3-5, 20*^ Bacteriophage T7 RNAP also bends the downstream duplex in the process of DNA opening^*21-24*^.

CryoEM studies indicate that ATP-independent promoter opening by eukaryotic RNA polymerases involves a similar set of conformational changes. In the assembly of the Pol II pre-initiation complex (Pol II PIC), the upstream DNA is bent by 90° by binding to the TATA element of transcription factors TBP (TATA-binding protein) together with TFIIA, TFIIB, and TAFs (TBP-associated factors).^*25-30*^ After Pol II PIC assembly, cryoEM reveals three closed complexes, designated CC1, CC2, and CC^dist^, which appear to be on-pathway intermediates in OC formation^*31*^. In CC1, the downstream promoter is positioned above the closed clamp of RNAP. In CC2 the clamp domain is open and in CC^dist^ the downstream duplex is bent into the open clamp. In the conversion of CC^dist^ to the final OC, the clamp closes and the DNA is opened^*31*^.

For *E. coli* RNAP, the isomerization of {CC} to OC is greatly facilitated by the presence of far-upstream DNA. For full-length (FL) λP_R_ promoter DNA, real-time HO footprinting of {CC} revealed interactions with RNAP up to at least position −82. ^*15*^ Upstream truncation of λP_R_ and lacUV5 promoters at positions between −63 (UT-63) and −42 (UT-42) reduces the isomerization rate constant k_isom_ (Mechanism for conversion of {CC} to OC by one to two orders of magnitude. ^*16, 17*^ For the UT-47 λP_R_ truncation variant, DNase footprinting of {CC} early in the time course of OC formation revealed downstream boundaries of partial protection of the template and nontemplate strands of the downstream duplex at +2 and +7, compared to +20 for {CC} at FL λP_R_.^*15*^ This indicates that the {CC} population distribution is much less advanced for UT-47 than for FL λP_R_. To explain these profound differences in OC-formation kinetics and {CC} footprints for FL and UT-47 λP_R_, we previously proposed that bending and wrapping of FL λP_R_ upstream DNA in an earlier closed complex (I_1E_) facilitates bending of the downstream duplex into the clamp/cleft in the most advanced member of the {CC} ensemble (I_1L_), in which the duplex is poised to be opened by RNAP.^*4, 15*^ The absence of upstream wrapping in UT-47 was proposed to greatly disfavor conversion of I_1E_ to I_1L_, thereby reducing k_isom_.^*4*^ Tests of these proposals are provided by the FRET and PIFE kinetics studies reported here.

Equilibrium FRET studies on low temperature (2 °C; closed) and higher temperature (19 °C; open) complexes revealed that far-upstream DNA in both complexes is highly bent and wrapped on RNAP.^*32*^ DNA backbone footprinting of RNAP-λP_R_ complexes revealed far-upstream protection to approximately −82 ({CC})^*15*^ and −65 (OC)^*15, 33*^, interpreted as upstream bending and wrapping, and AFM compaction measurements indicated extensive upstream wrapping in OC.^*34*^ Upstream modulators of initiation that bend and in some cases wrap DNA like CAP^*35*^, IHF^*36-38*^ and HU ^*39-41*^ may therefore affect k_isom_ by altering the distribution of wrapped and unwrapped intermediates in the {CC} ensemble and/or the trajectory of bending and wrapping of upstream DNA on RNAP in these intermediates.

In addition to FRET kinetic studies of changes of DNA end-to-end distance from bending and wrapping on RNAP, we report the kinetics of Cy3 and Cy5 fluorescence enhancements (PIFE)^*42-44*^ induced by contacts between RNAP and these dyes at the −100 and +14 positions of promoter DNA. The PIFE kinetics complement the FRET kinetics by reporting on the development of these RNAP-promoter contacts in {CC} intermediates of OC formation. PIFE effects using Cy3 at positions near the TSS were previously used to study aspects of the kinetics of OC formation, initiation, and escape.^*45-47*^ Cy3 PIFE kinetic measurements were also used to study the cooperative interaction between CarD and RbpA transcription factor in OC formation for *Mycobacterium tuberculosis* σ^A^ RNAP holoenzyme.^*48-50*^

Here we use FRET and PIFE in kinetic assays of OC formation to investigate large DNA conformational changes and changes in interactions of upstream and downstream DNA with RNAP as the initial CC advances and converts to OC. Analysis of the FRET and PIFE kinetic data provides compelling evidence for a sequential, five step mechanism with four closed complexes (the initial closed complex (RP_C_) and three more advanced members of {CC} designated I_1E_, I_1M_, I_1L_) on the pathway to open complex formation.

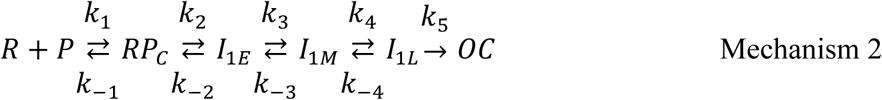

Illustrations of these intermediates, deduced from the research reported here, are provided in the graphical abstract and in Discussion.

Rate and equilibrium constants for reversible formation of the members of {CC} (RP_C_, I_1E_, I_1M_, I_1L_) are obtained, as well as the DNA opening rate constant k_open_ and intrinsic FRET and PIFE signatures of these closed intermediates to compare with those of OC and free P. FRET signatures of the intermediates provide information about the progression of upstream wrapping and downstream bending as {CC} advances. PIFE signatures of the intermediates provide information about the strength of contacts between RNAP and far-upstream (−100) and downstream (+14) regions of duplex DNA in early vs late CC and OC. The placement of the downstream duplex and the status of the RNAP clamp in the different CC intermediates deduced from the FRET and PIFE studies reported here relate well to cryoEM observations and proposals for pol II^*31*^ and E. coli RNAP intermediates^*46, 51*^. We also predict the time-evolution of the populations of individual early and late CC and the final OC. Rate and equilibrium constants obtained from FRET and PIFE analyses of OC formation at the λP_R_ promoter are compared with previous determinations of the composite quantities K_CC_ and k_isom_ for λP_R_ by filter binding assays.

These studies provide novel insights into the time-evolution of the {CC} ensemble and how wrapping of upstream DNA facilitates isomerization in OC formation. They provide strong support for the proposal that increases in the isomerization rate constant k_isom_ when upstream DNA or upstream-binding transcription factors are present, as well as from changes in promoter sequence, result from changes in the {CC} population distribution to favor the advanced closed complex I_1L_, in which the start site region of duplex promoter DNA is poised for DNA-opening in the RNAP clamp/cleft, and not primarily from increases in the rate constant of the subsequent DNA-opening step.^*4, 52, 53*^

## Materials and Methods

### Buffers

Storage Buffer (SB) for RNA polymerase holoenzyme is 50% glycerol (v/v), 10 mM Tris base (pH 7.5 at 4 °C), 0.1 M NaCl, 0.1 mM DTT, and 0.1 mM Na_2_EDTA. Fluorescence buffer (FB) for FRET and PIFE kinetics experiments is 40.2 mM Tris, pH 8, 2 mM NaCl, 0.12 M KCl, 10mM MgCl_2_, 2 μM DTT, 2 mM Na_2_EDTA, 0.05 mg/ml BSA, 0-2% glycerol and 0.02% Tween. Permanganate footprinting buffer (PFB) is 40 mM Tris (pH 8 at 19 °C), 10 mM MgCl_2_, 0.12 M NaCl and 100 μg/ml BSA. Urea loading buffer (ULB), used to resuspend footprinting samples, is 8 M urea, 0.5 X TBE (45mM Tris-borate and 1 mM Na_2_EDTA), 0.05% xylene cyanol (w/v) and 0.05% bromophenol blue (w/v).

### Preparation of E. coli RNA Polymerase (RNAP) Holoenzyme and Labeled λP_R_ Promoter DNA

RNAP core enzyme was overexpressed and purified as described previously, using *E. coli* BL21(λDE3) transformed with pVS10.^*32, 54*^ σ^70^ was overexpressed and purified using *E. coli* M5219 transformed with plasmid pMRG8, as described previously^*32*^. RNAP core enzyme and σ^70^ were incubated at a 1:2 molar ratio for 1 hour at 37°C in SB to reconstitute RNAP holoenzyme, then stored at −20°C until use. Other experiments were performed with preparations of WT RNAP holoenzyme.^*53, 55*^ Holoenzyme activities, determined from promoter binding assays at high promoter concentration,^*6*^ were in the range 50-90%. No significant differences between different RNAP preparations were observed in experiments reported here.

Single-dye-labeled λP_R_ fragments [Cy3 (−100), Cy5 (−100), Cy3 (+14) and Cy5 (+14)] for PIFE experiments and two-dye-labeled λP_R_ promoter DNA fragments [Cy3 (+14) Cy5 (−100) and Cy3 (−100) Cy5 (+14)] for FRET experiments were prepared by PCR as described previously using dye-labeled primers. ^*32*^ ^32^P-labeled λP_R_ promoter DNA fragments for MnO_4_^−^footprinting experiments were prepared by PCR as described previously.^*52, 55*^ Sequences and lengths of the different primers and DNA constructs are described in the supporting information (Tables S1-2).

### Fast MnO_4_^−^ Footprinting

RNAP and λP_R_ promoter DNA (−59 to +34) in PFB were independently loaded into a KinTek Corporation RQF-3 Rapid Chemical Quench-Flow instrument at 19°C. The solutions of RNAP and promoter DNA were mixed and held for varying times before mixing with 66.7 mM NaMnO_4_ for 50 ms, previously determined to be a concentration and time adequate for accurate detection of the rate of OC formation^*52*^. Samples were expelled, quenched with 500 µl ethanol, and immediately precipitated with ethanol and washed. Modified fragments were cleaved by 1 M piperidine at 90°C. Reactions were evaporated and resuspended in ULB and resolved on 8% acrylamide sequencing gels.

Each lane and reactive band of a given MnO_4_^−^ footprint was boxed and the total intensity was quantified using ImaqeQuant TL. The fraction of promoter DNA modified at a given position was determined by dividing the intensity of each uncut and reactive band by the total intensities of all uncut and reactive bands within a lane, and background was subtracted to determine corrected intensity. These corrected intensities were plotted vs time and fit to a single exponential time course. Corrected intensities were normalized by the appropriate fitted plateau intensity and plotted as normalized reactivity vs time.

### Kinetics of Open Complex Formation from Free RNAP and Promoter DNA by Stopped-Flow Fluorescence (FRET, PIFE)

Fluorescence-detected kinetic experiments were performed at 19°C in a Kintek SF-300X stopped flow fluorimeter (Kintek Corporation, PA) equipped with a 150 watt Hg-Xe lamp (Hamamatsu, Japan) by rapid-mixing of equal volumes (20 µL) of dye-labeled promoter DNA and RNAP stock solutions. Each mixing of aliquots of the same stock solutions is called a “shot”. Previous equilibrium FRET results on OC^*32*^ were obtained at 19°C because the fluorescence of cyanine dyes decreases strongly with increasing temperature^*32, 56*^. This temperature is used for kinetics experiments as well because at 19°C the equilibrium constant for forming the CC ensemble from promoter DNA is near-maximal, the rate of isomerization of the CC ensemble to OC is sufficiently slow to separate this kinetic phase from the earlier CC phase, and the final OC is sufficiently stable that its formation is irreversible.^*5, 57*^

Stocks of dye-labeled promoter DNA and RNAP were prepared at twice the desired final concentration in FB. Most experiments were performed at either 1:1 or 0.5:1 mole ratio of RNAP to promoter DNA (50 nM final DNA concentration), in order to avoid a competitive binding mode observed previously on this promoter DNA fragment in RNAP excess^*32*^. Control experiments (5-6 shots) were performed by mixing a DNA solution in FB with no RNAP to verify that the fluorescence signal was time-independent, without photobleaching or large instrumental drift. DNA and RNAP solutions were loaded in the stopped flow syringes and incubated for at least 10 minutes at 19°C before being mixed in a series of shots.

Dye-labeled λP_R_ promoter constructs were excited in the observation cell at wavelengths of 515nm (Cy3) or 610nm (Cy5) for single dye Cy3/Cy5 PIFE experiments. In FRET experiments, Cy3 dye (FRET donor) was excited at 515 nm wavelength and fluorescence emission of Cy3 and Cy5 (FRET acceptor) were monitored as a function of time. Fluorescence emission signals were monitored using a 565-625 nm band pass filter for Cy3 and a 660 nm long-pass filter for Cy5 (Omega Optical, Brattleboro, VT). A monochromator excitation slit width of 1.56 mm or 3.14 mm was used. Fluorescence intensity was monitored from 10ms to 400s with 600 data points that typically were uniformly distributed on a log time scale.

In RNAP-DNA mixing experiments the initial 4 shots were typically discarded and the next 5 - 15 shots were collected and analyzed. FRET data collected from each shot were normalized as described in supplemental and then averaged to reduce noise at earlier time points for further analysis. A total of 15 FRET experiments (each ~10 shots) were performed with Cy5 acceptor at −100, and another 12 experiments were performed with Cy5 at +14. In PIFE experiments, individual shots were also normalized and averaged. For comparative analysis of PIFE effects at −100 and +14, fluorescence increases relative to the initial (10 ms) signal were determined for each shot and averaged. A total of 34 PIFE experiments were performed: 15 monitoring +14 PIFE (10 with Cy3+14, 5 with Cy5+14) and 19 monitoring −100 PIFE (11 with Cy3-100, 8 with Cy5-100). Different experiments used independent dilutions (and in some cases independent preparations) of RNAP and DNA solutions. Analyses of these data in terms of a sum of exponentials and in terms of the proposed five-step mechanism are described in Supplemental Methods. Curves shown in Figs. 3 and 4 in Results are averages of series of normalized shots obtained in single experiments, selected as high S/N examples but otherwise representative of the full sets of independent experiments.

### Salt-Upshift Dissociation Kinetic Assays Monitored By FRET and +14 PIFE

Dye-labeled OC, prepared as described above, were rapidly mixed with KCl (50nM final OC concentration; 0.4 M final KCl concentration) in FB in the stopped-flow fluorimeter. Fluorescence kinetic data collected in each shot were normalized, and results from series of 10 or more shots were averaged as described above. FRET and PIFE curves shown in Results are representative of 4 independent FRET experiments and 5 independent PIFE experiments.

### Illustrating the Bending and Wrapping of Promoter DNA in Open Complex Formation

The PYMOL Molecular Graphics System, Version 2.3.2 (Schrödinger, LLC) was used to replace part or all of the DNA in the crystal structure of a transcription initiation complex (4YLN) by segments of DNA duplex (−100 to +14) to illustrate a plausible path for bending and wrapping of upstream DNA and bending of downstream DNA in the closed intermediates in OC formation as well as the final OC. These proposals for the approximate location of upstream and downstream duplex DNA in the bent-wrapped CC intermediates were developed to be semi-quantitatively consistent with the FRET distances and PIFE contacts determined in this research.

## Results

### Fast Permanganate Footprinting Kinetics of OC Formation

To test whether opening of the λP_R_ initiation bubble occurs as a single kinetic step and to validate the interpretation of the kinetics of open complex formation obtained previously from filter-binding data using Mechanism 1, fast permanganate (MnO_4_^−^) footprinting studies were used to determine rates of opening individual thymines in the λP_R_ open region (−11 to +2) in open complex formation at 19°C. Results are shown in Fig. 1. Fast MnO_4_^−^ footprinting of thymines in the −10 and start site regions of the promoter was previously performed to characterize the kinetics of open complex formation at the T7A1 promoter^*13*^ and to quantify the size of the bubble and the extent of reactivity of individual thymines in the unstable intermediate open complex (I_2_) formed in the DNA-opening step at the λP_R_ promoter. ^*52*^ Although MnO_4_^−^ reaction kinetics are moderately slow, requiring a reaction time of 150 ms even at high MnO_4_^−^ concentration, this is fast relative to the time scale of the CC-to-OC isomerization at 19°C (1/k_isom_ ~ 70 s). ^*5*^ Since only stable OC are detected, the MnO_4_^−^ kinetic assay is expected to be equivalent to the filter binding kinetic assay in which brief exposure of each sample to a competitor before assaying for binding eliminates contributions to the binding assay from any short-lived (closed or nonpromoter) complexes that were present.

**Figure 1.**
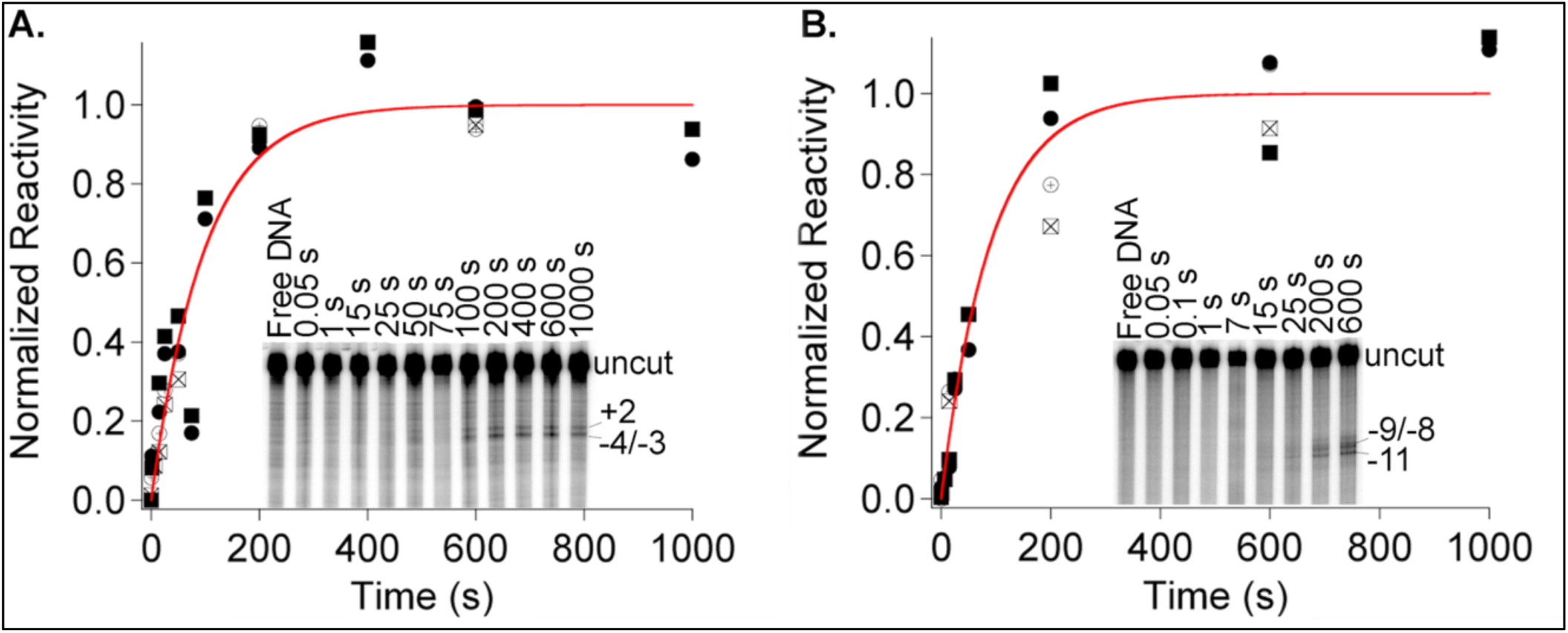
Kinetics of OC Formation Monitored by MnO_4_^−^ Reactivity. Fast (0.15 s) MnO_4_^−^ snapshots^*52*^ monitor the time course of opening individual thymines in OC formation after mixing excess RNAP (55 nM) with λP_R_ promoter DNA (0.3 nM) at 19 °C. Representative gels are shown as insets. Kinetics of development of MnO_4_^−^ reactivity are plotted for thymines on the nt strand (+2 (⊕, ●) and −4/-3 (⊠, ◼) in panel **A** and for the t strand (−9/-8 (⊕, ●) and −11 (⊠, ◼) in panel **B**. Rate constants k_obs_ for OC formation from these fits are the same within uncertainty (template strand k_obs_ = 0.010 ± 0.002 s^-1^; non-template strand k_obs_ = 0.011 ± 0.002 s^-1^). Times indicated are times after mixing RNAP with DNA at which the 0.15 s MnO_4_^−^ snapshot was initiated. Gels were quantified and results normalized as described in Materials and Methods.

Gel lanes in the insets in Fig. 1 show the kinetics of development of MnO_4_^−^ reactivity in the region of the initiation bubble on non-template (nt, panel A) and template (t, panel B) strands. In the first ~10 s, these MnO_4_^−^ snapshots detect no reactive (open) thymines (Fig S1), indicating that only CC complexes are present. MnO_4_^−^ reactivity of all observable thymines (−4/-3 and +2 on the nt strand; −11 and −9/-8 on the t strand) develops on a much slower timescale, becoming visible in the gel lanes only after 15 s and increasing to a plateau at times greater than 200 s. Single exponential global fits to the data for each strand (Figs. 1, S1) yield values of k_obs_, the observed first order rate constant for the formation of MnO_4_^−^-detected complexes at 55 nM excess RNAP, which are the same within the uncertainty (10-20%) for all thymines detected on both template and nontemplate strands.

Rate constants k_obs_ obtained from these MnO_4_^−^ footprinting assays at 19 °C are compared with those obtained previously by filter binding^*5*^ at 20 °C in a plot vs [RNAP] in Figure 2A, and were fit to the expected hyperbolic functional form:^*1-7*^

**Figure 2.**
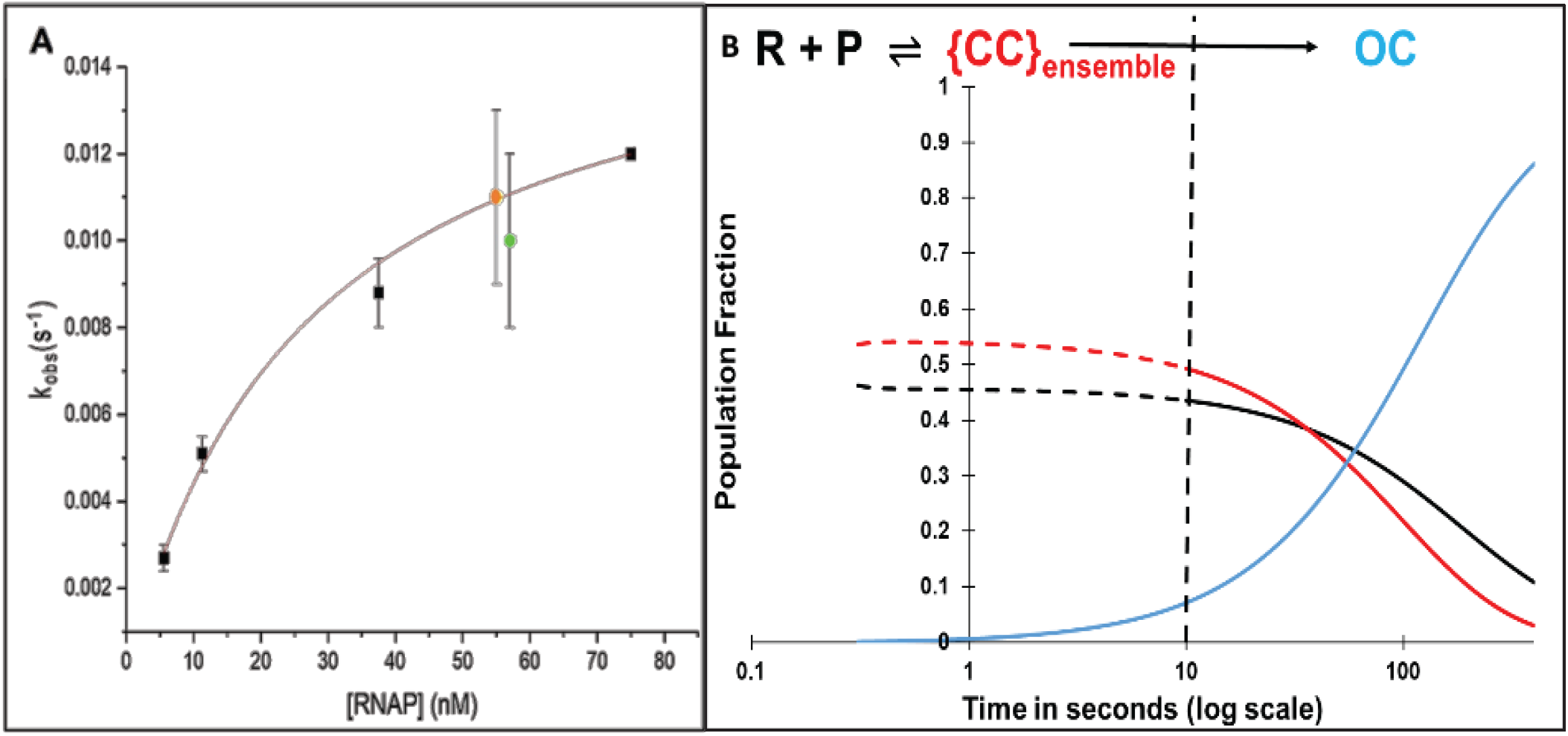
Panel A: Dependence of Rate Constant k_obs_ for λP_R_ OC Formation on RNAP Concentration in RNAP Excess. First order rate constants k_obs_ (Eq. 1) for OC formation in excess RNAP are plotted as a function of RNAP concentration. ◼: k_obs_ values from filter binding assays at 20°C.^4^ ●, ●: k_obs_ values from MnO_4_^*-*^ footprinting of template and non-template strands, respectively at 19 °C (Fig 1). A fit of these data to hyperbolic Eq. 1 gives a composite {CC} binding constant K_{CC}_ = (5 ± 1) × 10^7^ M^-1^ and an isomerization rate constant k_isom_ = 0.014 ± 0.003 s^-1^ for conversion of {CC} to OC. **Panel B: Simulated Time Evolution of {CC} and OC Formation for FRET/PIFE Conditions (50 nM λP**_**R**_ **Promoter and RNAP).** Fractional populations of open complexes (OC) —, closed complexes {CC} —, and free promoter DNA —, predicted from K_CC_ and k_isom_ as a function of time (log scale) assuming that the {CC} population equilibrates with free promoter DNA by 10 s (_ _ _and _ _ _). The dashed vertical line at 10 s marks the onset of OC formation from the equilibrium mixture of {CC} and free promoter DNA determined from this simulation.

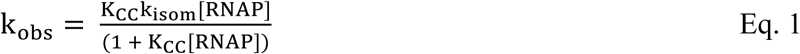

where K_CC_ is the equilibrium constant for forming the CC ensemble from RNAP and promoter DNA, and k_isom_ is the CC-to-OC isomerization rate constant. Values of K_CC_ and k_isom_ obtained from this analysis (see Fig. 2 caption) agree within the uncertainty with those reported previously.^*5*^

### Predicted Time Evolution of CC and OC Populations at RNAP and Promoter Concentrations of FRET and PIFE Experiments

The kinetic results (K_CC_, k_isom_) from the analysis in Fig. 2A, obtained in RNAP excess at a low promoter concentration (0.3 nM) allow prediction of two key aspects of the time course of open complex formation for the very different conditions of the fluorescence experiments (1:1 or limiting (e.g. 0.5:1) RNAP; high (50 nM) DNA concentrations). The extent of conversion of P to {CC} in the rapidly reversible first step of Mechanism 1 is predicted by K_CC_ and the kinetics of conversion of this rapidly-reversible mixture of CC and P to OC is predicted by k_isom_. These predictions, given in Fig. 2B for 50 nM RNAP and promoter DNA on a logarithmic time scale, show that significant OC formation does not occur in the fluorescence kinetics experiments until approximately 10 s after mixing.

Hence the kinetics of OC formation are predicted to exhibit *two phases*. The first phase (0 – 10 s after mixing for the conditions simulated) is the step-wise formation of the ensemble of closed complex intermediates ({CC}) from free promoter DNA. Because the kinetics of OC formation in filter binding and MnO_4_^−^ reactivity assays that detect only OC are single-exponential in excess RNAP, formation of the {CC} ensemble (Mechanism 1) and hence the individual steps of {CC} formation (Mechanism 2) are rapidly reversible^*58*^. Equilibrium between {CC} and P is therefore established in the first 10 s of the reaction, quantified by the equilibrium constant K_CC_.

At 10 s, Fig. 2B predicts that approximately half of total promoter DNA is bound in closed complexes ({CC}) with RNAP while the other half is free (unbound) P. This {CC} ensemble (and free P) convert to OC in the second kinetic phase. The kinetics of the steps of evolution of {CC} in the transient first phase cannot be obtained from studies of OC formation like those in Fig 1, but are determined by fitting of FRET and PIFE kinetic data for these dye positions to the five-step Mechanism 2

### FRET-Detected Kinetics of DNA Bending and Wrapping in Closed and Open Complex Formation

Previous FRET studies of equilibrium populations of closed (2°C) and open (19°C) complexes of RNAP and doubly-labeled (Cy3 and Cy5) λP_R_ promoter DNA revealed that upstream and downstream DNA regions are highly bent and upstream DNA is wrapped on RNAP in the low-temperature (2 °C) population of closed complexes and in the 19 °C open complex, reducing the distance between −100 and +14 positions (numbered by convention relative to the +1 start site) from >300 Å before binding RNAP to 60-70 Å in these complexes. To determine the time course and mechanism of these DNA deformations, FRET kinetics experiments were performed at 19° for Cy3(−100)Cy5(+14) and Cy3(+14)Cy5(−100). Representative FRET acceptor time-courses for both promoter fragments, plotted on a logarithmic time scale, are shown in Fig. 3.

**Figure 3.**
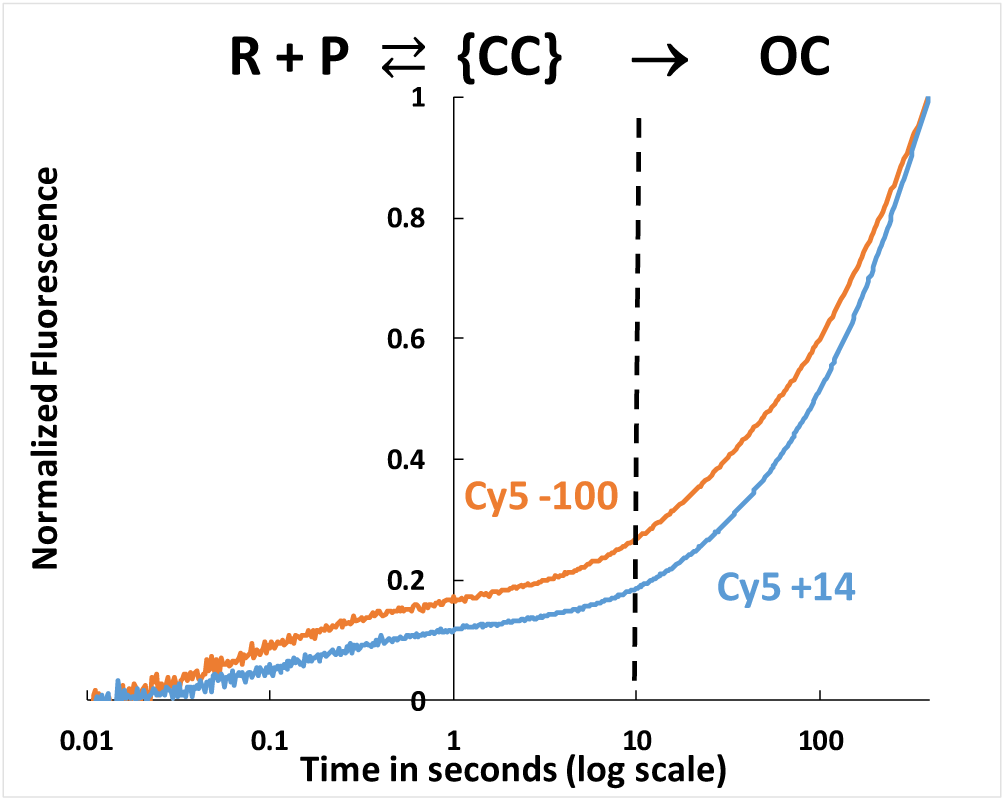
FRET-detected Bending and Wrapping of Promoter DNA by RNAP as {CC} Ensemble Advances and Forms the Stable OC. Time course (log scale) of normalized FRET acceptor (Cy5) emission intensity after mixing Cy3-Cy5 dye-labelled λP_R_ DNA (50 nM final) with *E. coli* RNAP (50 nM final) at 19°C and exciting FRET donor (Cy3) at 515nm. Both dye orientations are shown: Cy5(+14) Cy3(−100) —, Cy3(+14) Cy5(−100) —. The vertical dashed line at 10 s corresponds to the onset of OC formation at these conditions.

The FRET acceptor kinetics span a time range of more than four orders of magnitude, from ~20 ms to at least 400 s. As predicted in Fig. 2B, two kinetic phases are observed. The faster phase, accounting for approximately 15% of the total FRET increase, occurs in the first 10 s after mixing, and the slower second phase develops from 10 s to 400 s. By comparison with the simulations of Fig. 2B, these two kinetic phases clearly correspond to reversible formation of the {CC} ensemble from reactants, followed by conversion of this {CC} ensemble and free promoter DNA to OC. The FRET inflection point in Fig. 3 in the time range 1-10 s, when ~50% of promoter DNA has been converted to the {CC} ensemble, is only ~15% of the long-time (400 s) value, at which ~80% of promoter DNA is in an OC (Fig 2B). This indicates that the {CC} ensemble is on average less bent/wrapped, with a larger dye-dye distance, than the final OC. The two high S/N experiments plotted in Fig. 3 are otherwise representative of the full set of 27 FRET experiments. In the time range from 0.5 s to 200 s the normalized FRET signal for Cy5+14 is generally smaller than for Cy5-100.

### RNAP-Induced Fluorescence Enhancements (PIFE) at Both −100 and +14 Exhibit Similar Rates to FRET Acceptor Increases but Decrease Late in {CC}. Phase and in OC Formation

Single-dye kinetics experiments (Fig. 4) show large increases in Cy3 and Cy5 fluorescence (PIFE effects) in the time range 10 ms to 1 s at both −100 and +14 positions of λP_R_ promoter DNA. −100 and +14 PIFE signals in Fig. 4 both develop on the same time scale as the FRET acceptor signal (t > 10 ms; Fig. 3). These +14 and −100 PIFE effects provide kinetic information regarding the involvement of downstream and upstream DNA in steps of OC formation. At each position, Fig. 4 shows that normalized PIFE effects of Cy3 and Cy5 are similar. PIFE effects for both dyes at both positions exhibit two kinetic phases, as in FRET assays (Fig. 3). In all four cases, PIFE effects are first detectable approximately 10-20 ms after mixing and increase to a maximum near the end of the {CC} phase. For +14 PIFE, a broad maximum in the time range 2 – 20 s is observed (Fig. 4A). For −100 PIFE the maximum is sharper and occurs earlier (1-3 s). For t > 3 s (−100 PIFE) and t > 20 s (+14 PIFE). PIFE signals decrease as the {CC} ensemble is completed and conversion to the stable OC begins, indicating that intrinsic −100 and +14 PIFE signal intensities of {CC} intermediates like I_1E_ and I_1M_ exceed those of the stable OC (see Discussion).

**Figure 4.**
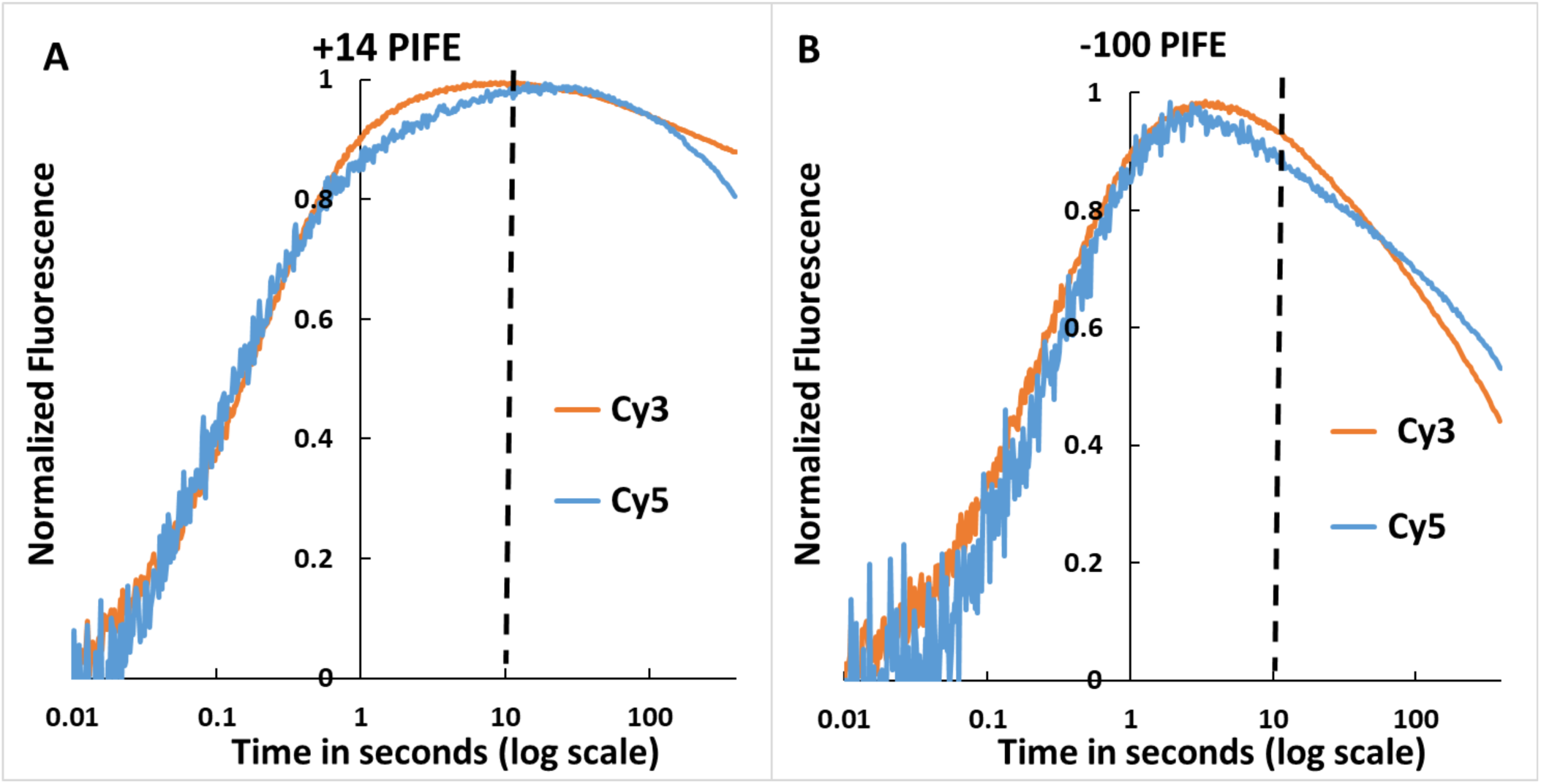
OC Formation Monitored by Single-Dye Cy3, Cy5 Fluorescence (PIFE) at Downstream (+14) and Far-Upstream (−100) Positions of λP_R_ Promoter DNA. Representative time courses (log scale) of normalized single-dye fluorescence (PIFE) of Cy3- or Cy5-labeled λP_R_ DNA after mixing with RNAP (final concentrations 50 nM RNAP and λP_R_ DNA) at 19 °C. +14 PIFE (**Panel A)** and −100 PIFE (**Panel B**) from Cy3 — and Cy5 —. Cy3 was excited at 515 nm and Cy5 at 610 nm. The vertical dashed line at 10 s is the predicted onset of OC formation (see Fig. 2B).

Position +14 is ~25 bp downstream of the footprint boundary for RP_c_ (typically −5) but within the +20 downstream boundary of the footprint observed for more advanced CC and OC.^*3, 4*^ Position −100 is 20-40 bp upstream of footprint boundaries of all RNAP-λP_R_ promoter CC and OC investigated. Nevertheless, the PIFE results demonstrate that RNAP must contact^*44*^ these dye positions on promoter DNA in the step(s) that advance the {CC} ensemble by bending and wrapping upstream and downstream promoter DNA on RNAP. The Discussion considers the implications of these contacts for the operation of the RNAP-promoter biophysical machinery that uses binding free energy to bend the downstream duplex onto the top of the β pincer and then into the open clamp, before clamp-closing and opening of 13 base pairs including the transcription start site.

### Unwrapping in Dissociation of OC by a Salt-Upshift

Rapid salt upshifts were used to investigate unwrapping in OC dissociation. Mixing of a stable RNAP-λP_R_ promoter OC with high KCl or urea rapidly destabilizes it, forming the unstable but [KCl]-insensitive open-promoter intermediate I_2_, which undergoes DNA closing on a 1 s time scale^*52, 53, 59, 60*^. Upshifts of doubly- and singly-labeled OC to 0.4 M KCl at 19°C result in rapid reductions in Cy5+14 acceptor FRET and Cy3 +14 PIFE. In Fig. 5 the kinetics of these fluorescence changes are compared with one another and with simulations of the two-step irreversible conversion of the stable OC at 19 oC to closed promoter complexes via the unstable open-promoter intermediate I_2_ and the DNA-closing step ^*52, 53, 59, 60*^. Figure 5 shows that the FRET and PIFE fluorescence changes occur prior to the DNA closing step of high-salt-induced dissociation. The FRET decrease occurs faster than the reduction in +14 PIFE. Both are slower than the conversion of RP_o_ to I_2_ but faster than the conversion of I_2_ to closed-promoter complexes. The finding that the FRET decrease occurs faster than the reduction in +14 PIFE indicates that KCl-driven unwrapping of upstream DNA occurs before the release of downstream contacts. This sequence of steps in response to a large KCl-upshift differs from the mechanism of OC formation at lower KCl, where upstream DNA wrapping and formation of +14 contacts occur together as the {CC} ensemble advances.

**Figure 5.**
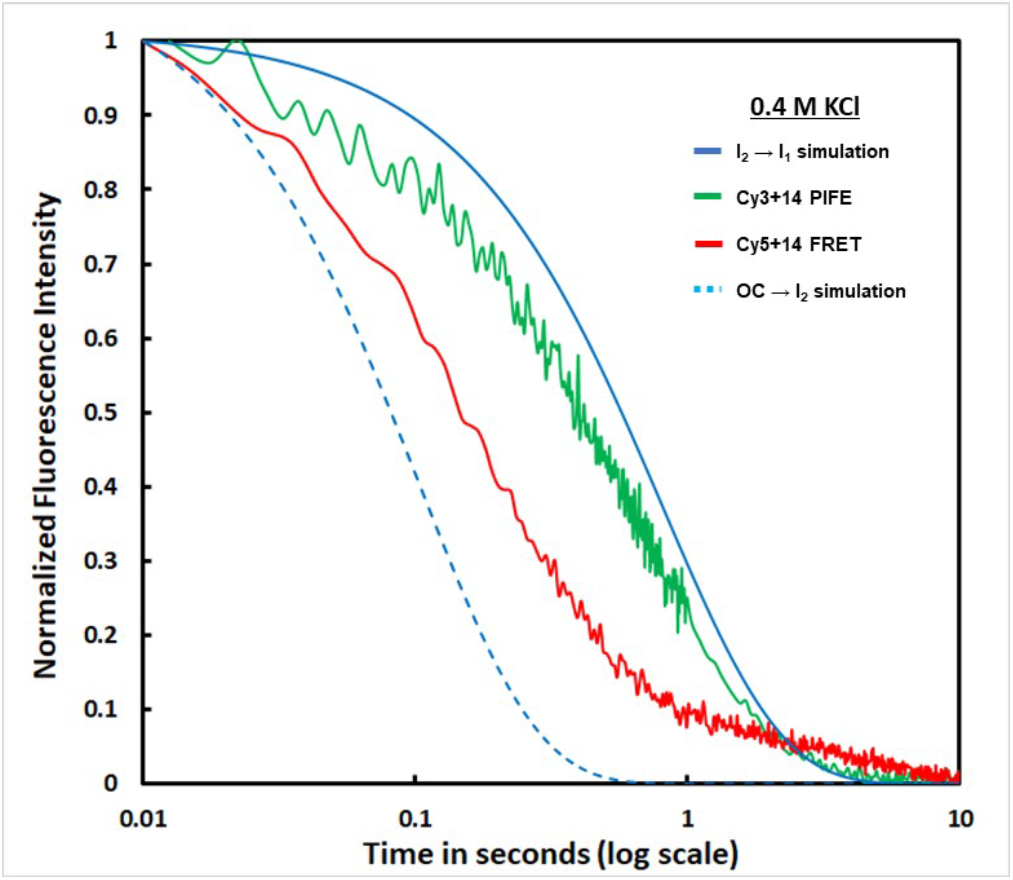
Unwrapping (detected by FRET) and Release of Downstream Contacts (detected by +14 PIFE) in OC Dissociation after an Upshift to 0.4 M KCl at 19 °C. Representative time courses (log scale) of reductions in Cy5+14 acceptor FRET — and Cy3+14 PIFE — after destabilizing the 19 °C OC with 0.4 M KCl. Blue curves are simulations based on 0.4 M KCl rate constants, interpolated to 19 °C from results at 10 °C and 37 °C.^*59*^ These predict the time course of conversion of the initial OC to the unstable open intermediate I_2_ (OC → I_2_, rate constant k_-3_ = ~9.7 s^-1^; _ _) and subsequent DNA-closing, designated as I_2_ → I_1_ with rate constant k_-2_ = ~1.2 s^-1^; —.

## Analysis and Discussion

### A Five-Step Mechanism of OC Formation with a Key Intermediate CC (I_1M_) Not Previously Observed by DNA Footprinting

OC formation, studied in RNAP excess by MnO_4_^*-*^ footprinting and filter binding assays that only detect the stable OC (e.g. Fig. 1), occurs over two decades in time and exhibits single-exponential kinetics (Fig. S1). The RNAP concentration dependence of the kinetics is well described by two-step Mechanism 1 (parameters K_CC_, k_isom_). The single-exponential kinetics of OC formation in these experiments show that formation of the {CC} ensemble equilibrates rapidly (characterized by K_CC_) on the time scale of the conversion of {CC} to OC (characterized by k_isom_). Because of {CC} rapid equilibrium, no information is obtained in these kinetic studies about the number of CC intermediates or the kinetics of their interconversions.

On the other hand, four exponentials with rate constants 1/*τ*_*i,obs*_ that span four decades (~ 15 s^-1^ to ~ 0.015 s^-1^; Table S3) are needed to fit the FRET (Fig. 3) and PIFE (Fig. 4) kinetic data for OC formation, as described in SI. This indicates the presence of significant populations of at least three intermediates in the {CC} ensemble with FRET and/or PIFE fluorescence signals which differ from free promoter DNA, from one another and from the final OC. Because the {CC} ensemble is in rapid equilibrium with reactants on the time scale of its isomerization to OC, each step of advancing this ensemble also rapidly equilibrates on the time scale of the next forward step (i.e. k_-1_ > k_2_, k_-2_ > k_3_, etc.). Consequently, the three rate constants 1/*τ*_1,*obs*_, 1/*τ*_2,*obs*_ and 1/*τ*_3,*obs*_ in the four-exponential fit can be interpreted as decay-to equilibrium rate constants^*61*^ for formation of three observable CC (designated I_1E_, I_1M_, I_1L_; see SI Eqs. S3-S7). We deduce that rapid reversible formation of the initial specific CC (RP_C_) from reactants is not directly detected by FRET, nor by −100 or +14 PIFE, because promoter DNA is not sufficiently bent in RP_C_ to give FRET and because contacts between RNAP and the promoter in RP_C_ do not extend to −100 or +14. The final (fourth) exponential is interpreted as irreversible conversion of the most advanced closed intermediate (I_1L_) to the stable 19 °C OC in several steps, which are not separable because they follow the rate-determining DNA opening step (I_1L_ → I_2_, with rate constant k_5_).

Given the need for four exponentials to fit these data, it is not surprising that a five-step mechanism (Mechanism 2) including RP_C_, I_1E_, I_1M_, and I_1L_ is needed to obtain good fits to all the FRET acceptor and PIFE kinetic data using a common set of rate constants (see below). FRET and +14PIFE kinetic data can be adequately fit using a four step mechanism, but significantly different sets of rate constants are obtained for FRET and PIFE datasets. −100 PIFE kinetic data are not adequately fit by a four step mechanism.

Fits of the 5-step mechanism to the FRET and PIFE kinetic data of Figs. 3 and 4 are shown in Panels A-C of Fig. 6, using the best-fit amplitudes and rate constants for these individual data sets listed in Table S4. Clearly Mechanism 2 fits the experimental data with only minor deviations (Table S4). For comparison, Panels A-C of Fig. S2 show fits to the same FRET and PIFE time-courses using average FRET and PIFE signal amplitudes (Table S5) and average rate constants and obtained from fitting all 61 data sets to Mechanism 2. Use of these average amplitudes and rate constants also provides good fits with approximately two-fold larger deviations (Table S5).

**Figure 6.**
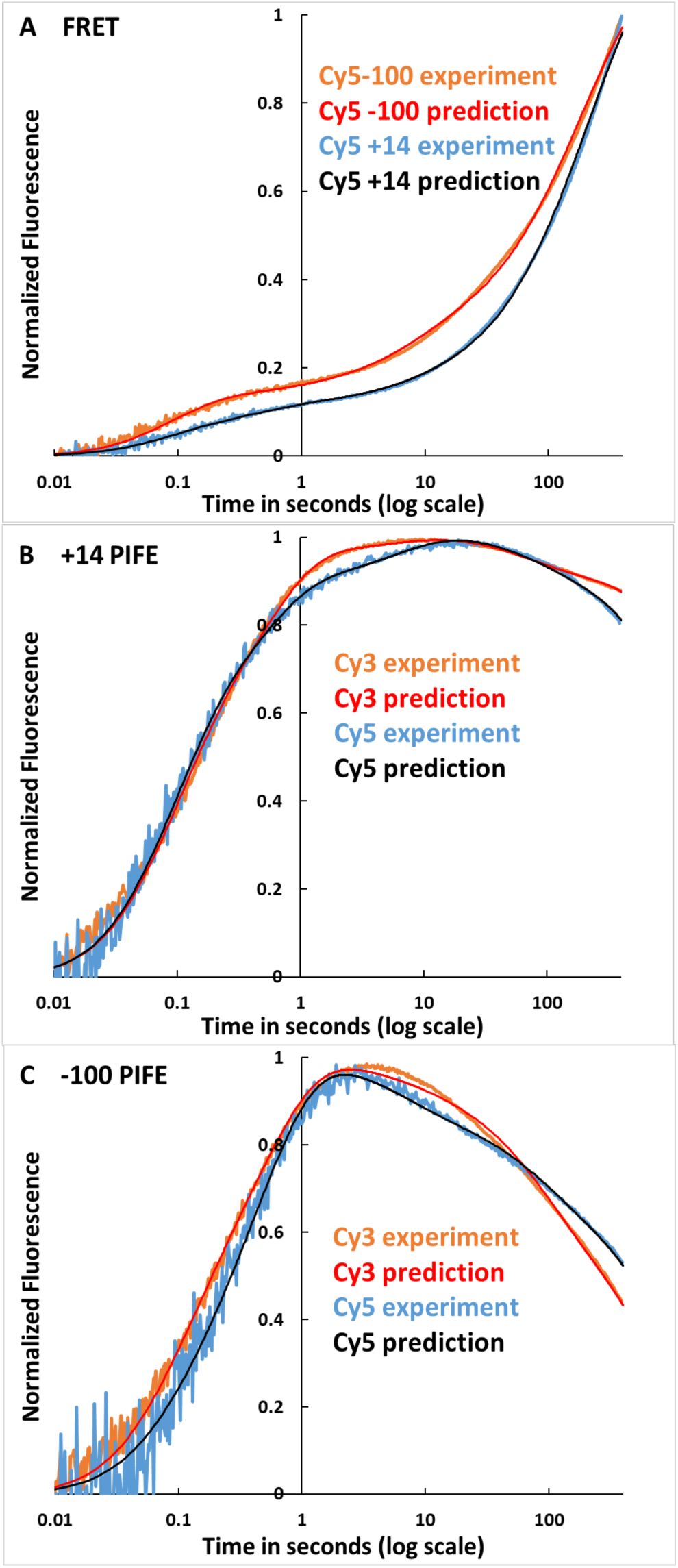
Comparison of Predicted and Observed FRET and PIFE Kinetics of OC Formation at λP_R_ Promoter. Panel A: FRET data (Fig. 3) for Cy5-100 — and Cy5+14 —. Panel B: +14 PIFE data (Fig 4A) for Cy3 — and Cy5 —. Panel C: −100 PIFE data (Fig. 4B) for Cy3 — and Cy5 —. Predictions use rate constants and amplitudes in Table S4, obtained from fitting these data sets to Eq. S7 for Mechanism 2. For comparison, fits in SI Fig. S2 use average rate constants (Table 3) and amplitudes (Table S5) for the entire data sets.

The following sections discuss the average FRET acceptor and PIFE amplitudes of the intermediate CC and OC in 5-step Mechanism 2, their implications for the large-scale conformational changes in promoter DNA and RNAP in these steps, the average rate and equilibrium constants of these steps, the simulated time-course of OC formation and the free energy vs progress diagram for this process.

### FRET Evidence that Promoter DNA is Wrapped on RNAP in All Three I_1_ Intermediates and that Far-Upstream (−100) and Downstream (+14) DNA are Farther Apart in I_1M_ than in I_1E_, I_1L_

Table 1 lists average Cy5 FRET acceptor signal amplitudes of {CC} intermediates for both Cy3(−100) Cy5(+14) and Cy3(+14)Cy5(−100), calculated from the average FRET acceptor fitting amplitudes in Table S5 and expressed relative to OC. Cy3 (FRET donor) fluorescence is dominated by PIFE and provides no FRET information. No Cy5+14 or Cy5-100 FRET is observed for the initial closed complex RP_C_, showing that the distance between - 100 and +14 exceeds the FRET detection limit (~100 Å). The three more advanced I_1_ intermediates in the {CC} ensemble all exhibit significant FRET between −100 and +14, indicating that these positions, more than 300 Å apart in free promoter DNA, are less than ~100 Å apart in these intermediates. FRET amplitudes of the I_1_ intermediates range from ~10-80% that of OC (Table 1). These observations are consistent with previous equilibrium FRET results which indicated that promoter DNA in the 2 °C {CC} ensemble is highly wrapped on RNAP, with a somewhat smaller FRET efficiency (i.e. larger −100/+14 distance) than the 19 °C OC^*32*^.

**Table 1.** Relative FRET and PIFE Fluorescence Signals of RNAP-Promoter Complexes.

|  | FRET Acceptor Amplitude (Between<br>-100 and +14, Relative to OC)<sup>a</sup> |  | +14 PIFE Intensity<br>(Relative to P)<sup>b</sup> |  | -100 PIFE Intensity<br>(Relative to P)<sup>b</sup> |  |
| --- | --- | --- | --- | --- | --- | --- |
|  | Cy5+14 | Cy5-100 | Cy3 | Cy5 | Cy3 | Cy5 |
| **RP<sub>c</sub>** | **0 ± 0.1** | **0 ± 0.1** | **1 ± 0.1** | **1 ± 0.1** | **1 ± 0.1** | **1 ± 0.1** |
| **I<sub>IE</sub>** | **0.34 ± 0.13** | **0.56 ± 0.26** | **1.66 ± 0.32** | **1.15 ± 0.08** | **1.20 ± 0.11** | **1.06 ± 0.03** |
| **I<sub>IM</sub>** | **0.13 ± 0.09** | **0.24 ± 0.13** | **1.53 ± 0.25** | **1.11 ± 0.06** | **1.19 ± 0.11** | **1.11 ± 0.04** |
| **I<sub>IL</sub>** | **0.71 ± 0.24** | **0.79 ± 0.15** | **1.42 ± 0.16** | **1.12 ± 0.06** | **1.04 ± 0.04** | **1.02 ± 0.02** |
| **OC** | **1** | **1** | **1.27 ± 0.10** | **1.07 ± 0.03** | **1.04 ± 0.04** | **1.02 ± 0.02** |

<sup>a</sup> Average values of $A_i/A_{OC}$ (Eq. S7) from fits of FRET kinetic data to Mechanism 2
<sup>b</sup> Average values of relative PIFE intensities ( $\frac{a_i}{a_P}$ ), calculated from fits of PIFE kinetic data to Mechanism 2 using Eq. S17.

These average FRET acceptor signal amplitudes of {CC} intermediates are analyzed to obtain relative FRET amplitudes, FRET efficiencies and distances between dyes at −100 and +14 in Table 2, using the previously-determined OC FRET efficiency (0.32).^*32*^ FRET efficiencies of I_1E_, I_1M_ and I_1L_ are ~45%, ~18% and ~75% that of OC. Use of the previously-determined −100/+14 OC distance (63 Å)^*32*^ as a reference yields a −100/+14 distance for I_1E_ of ~75 Å. The −100/+14 distance increases by ~14 Å in conversion of I_1E_ to I_1M_ before decreasing by ~21 Å in conversion of I_1M_ to I_1L_. The −100/ +14 distance in I_1L_ (~68 Å) is ~5 Å greater than in the stable OC. Details of this analysis are provided in SI.

**Table 2.** FRET-Determined Distances between −100 and +14 Probe Positions on λP_R_ Promoter DNA in CC Intermediates.

| **Table 2. FRET-Determined Distances between -100 and +14 Probe Positions on $\lambda P_R$ Promoter DNA in CC Intermediates** |  |  |  |
| --- | --- | --- | --- |
| **Species** | **FRET Acceptor Relative Amplitude<sup>a</sup>** | **FRET Efficiency** | **Distance between -100 and +14** |
| **$I_{IE}$** | **$0.45 \pm 0.27$** | **$0.14 \pm 0.08$** | **$75 \pm 9 \text{ \AA}$** |
| **$I_{IM}$** | **$0.18 \pm 0.16$** | **$0.059 \pm 0.052$** | **$89 \pm 14 \text{ \AA}$** |
| **$I_{IL}$** | **$0.75 \pm 0.30$** | **$0.24 \pm 0.09$** | **$68 \pm 6 \text{ \AA}$** |
| **OC** | **1** | **$0.32^b$** | **$63 \text{ \AA}^b$** |
| <sup>a</sup> Average of Cy5-100 and Cy5+14 results $\pm 1$ SD (Table 1) expressed relative to OC. | | | |
| <sup>b</sup> From Ref. 32. FRET efficiencies and dye-dye distances for $I_{IE}$, $I_{IM}$ and $I_{IL}$ are calculated by comparison to this OC value. | | | |

**Table 3.** Rate Constants and Equilibrium Constants for Formation of the Ensemble of Closed Complex Intermediates and DNA Opening at 19 °C.

|  | Kinetic Step | Rate Constants (± Standard Deviation) | Equilibrium Constant<sup>a</sup> |
| --- | --- | --- | --- |
| 1 | $R + P \rightleftharpoons RP_c$ | $k_1 = (3.0 \pm 0.2) \times 10^8 \text{ M}^{-1}\text{s}^{-1}$, $k_{-1} = 44 \pm 2 \text{ s}^{-1}$ | $K_1 = (6.8 \pm 0.5) \times 10^6 \text{ M}^{-1}$ |
| 2 | $RP_c \rightleftharpoons I_{1,E}$ | $k_2 = 13 \pm 1 \text{ s}^{-1}$, $k_{-2} = 7.2 \pm 0.4 \text{ s}^{-1}$ | $K_2 = 1.8 \pm 0.2$ |
| 3 | $I_{1,E} \rightleftharpoons I_{1,M}$ | $k_3 = 2.5 \pm 0.1 \text{ s}^{-1}$, $k_{-3} = 1.7 \pm 0.1 \text{ s}^{-1}$ | $K_3 = 1.5 \pm 0.1$ |
| 4 | $I_{1,M} \rightleftharpoons I_{1,L}$ | $k_4 = 0.10 \pm 0.01 \text{ s}^{-1}$, $k_{-4} = 0.10 \pm 0.01 \text{ s}^{-1}$ | $K_4 = 1.0 \pm 0.1$ |
| 5 | $I_{1,L} \rightarrow \text{OC}$ | $k_5 = 0.040 \pm 0.002 \text{ s}^{-1}$ <sup>b</sup> | |

<sup>a</sup> These four equilibrium constants yield a predicted $K_{\{CC\}} = (5.5 \pm 0.8) \times 10^7 \text{ M}^{-1}$ from Eqs. S21-22, in agreement with the experimental $K_{\{CC\}} = (5 \pm 1) \times 10^7 \text{ M}^{-1}$ (Fig. 2A).
<sup>b</sup> This $k_5$ , multiplied by the fraction of the equilibrated {CC} that is I<sub>1L</sub> calculated from K<sub>1</sub>, K<sub>2</sub>, K<sub>3</sub> and K<sub>4</sub> above (Eqs. S19-20), gives a predicted $k_{\text{isom}} = 0.013 \pm 0.001 \text{ s}^{-1}$ the same within uncertainty as the experimental $k_{\text{isom}} = 0.014 \pm 0.003 \text{ s}^{-1}$ (Fig. 2)

Although the uncertainties in these distances (Tables S5, 1, 2) are appreciable, comparable in some cases to the differences between the different I_1_ intermediates, the trend is unambiguous. These uncertainties result from the imperfect reproducibility of the data for each position of the Cy5 acceptor and from the marginally-significant difference in FRET signals for the two placements of the dye probes (Fig. 3). Separate analyses of Cy5-100 and Cy5+14 FRET acceptor data sets (Tables S4, S5) reveal the same progression of FRET fitting amplitudes for these intermediates, with I_1M_ exhibiting the smallest amplitude and therefore the greatest distance between −100 and +14 probe positions. The conversion of higher-FRET I_1E_ to low-FRET I_1M_ is the origin of the inflections in the Cy5+14 and Cy5-100 FRET acceptor time courses at about 1 s (Fig. 3).

These large changes in distance between −100 and +14 for the different intermediates, together with PIFE contact information at these positions, form the basis for a structural mechanism of OC formation, proposed below. They also provide a structural explanation of the very-large facilitation of isomerization in OC formation by upstream wrapping.

### PIFE Indicates that RNAP-Promoter Contacts at +14 and Especially at −100, Absent in RP_C_, Are Stronger in I_1_ Intermediates than In OC

Table 1 also summarizes the RNAP-induced fluorescence enhancements (PIFE) determined for the different CC intermediates and OC, expressed relative to free promoter DNA (P). These are calculated as described in SI (Eqs. S8-17) from the average PIFE fitting amplitudes (Table S5), which span a ~10-fold range. Several general trends are clear from Table 1. Of most significance, PIFE intensities at −100 and +14 are larger for I_1E_ and I_1M_ intermediates than for OC for both Cy3 and Cy5, while RP_C_ exhibits no detectable PIFE intensity. The −100 PIFE intensity of I_1L_ is small, comparable to the −100 PIFE intensity of OC, while the +14 PIFE intensity of I_1L_ is larger, comparable to the −100 PIFE intensities of I_1E_ and I_1M_. These rank orders of PIFE intensities indicate that relatively strong contacts^*44*^ of RNAP with both far-upstream (−100) and downstream (+14) DNA are made in converting RP_C_ to I_1E_, where −100/+14 FRET is also first observed. *These relatively strong contacts persist in I*_*1M*_, *demonstrating that the decrease in FRET in converting I*_*1E*_ *to I*_*1M*_ *does not result from partial unwrapping of upstream or downstream DNA.*

Contacts of RNAP with far upstream (−100) DNA appear to be largely disrupted in conversion of I_1M_ to I_1L_ and OC, while contacts of RNAP with downstream (+14) DNA appear similar in all three I_1_ intermediates and surprisingly are stronger in these intermediates than in the stable OC. In other trends, contacts of RNAP with +14 appear stronger than with −100, since the PIFE signal of each dye in each complex (I_1_ intermediates, stable OC) is stronger at +14 than at −100. Also, the effect of RNAP-DNA interaction on dye fluorescence is dye-specific, with larger Cy3 PIFE intensities than Cy5 PIFE intensities at each position.

### FRET/PIFE Insights into the Mechanism of Operation of the RNAP-Promoter Machine in OC Formation

#### a) Specific Contacts in RP_C_ Direct RNAP to Bend and Wrap Both Upstream and Downstream DNA in One Step to Form High-FRET/PIFE I_1E_

The development of −100/+14 FRET and −100 and +14 PIFE in conversion of RP_C_ to I_1E_ demonstrates that both upstream and downstream promoter DNA are bent and wrapped on RNAP in this step, making contacts between RNAP and both −100 and +14 positions of promoter DNA and reducing the −100/+14 distance to ~75 Å (Table 2). To interpret these concerted upstream and downstream effects, we propose that the contacts made in RP_c_ between RNAP and −35 and/or UP elements of promoter DNA initiate strong upstream bending, resulting from or accompanied by contraction/folding of the flexible tethers linking α-CTD to α-NTD, as discussed below. Upstream bending allows far-upstream wrapping, which, as proposed previously, allows the −100 position of the promoter to be contacted by a mobile downstream element (DME) or other region at the downstream end of the RNAP clamp^*3, 15*^. If only the upstream DNA but not the downstream DNA were bent and wrapped, the distance between −100 and +14 probe positions is estimated to be >130 Å, which is too large to exhibit FRET. Therefore bending and wrapping of upstream DNA must be accompanied by bending of the downstream duplex, presumably at the upstream end of the −10 region,^*5, 19, 20, 51, 62*^ reducing the distance between +14 and −100 positions on promoter DNA to ~75 Å to give the FRET efficiency for I1E calculated in Table 2. This downstream bending is not present in RPC, because downstream footprints of RP_C_ complexes end at −5 while that of I_1E_ extends to at least +2/+7.^*15*^ Hence the interactions of σ^70^ region 2 with the −10 region in RP_C_ are not sufficient by themselves to induce the large-scale bending of the downstream duplex observed in I_1E_. If other interactions of the downstream duplex with RNAP are involved in the conversion of RP_C_ to I_1E_, these might be with a RNAP element like the βSI1 sequence insertion or the NCD of σ^70^.

A plausible illustration of I_1E_ is provided in Fig. 7. A −100/+14 dye-dye distance of ~75 Å is obtained by bending the upstream duplex around the α subunits, wrapping it on the outside of the β’ clamp of RNAP, and bending the downstream duplex at the upstream end of the −10 region to bring +14 in contact with the top of the β pincer and sequence insertion βSI1. This model is similar in its placement of the downstream duplex to the CC1 pol II intermediate observed by cryoEM.^*31*^ A recent cryoEM study of allosteric regulation of RNAP by TraR finds that this protein also interacts with the top of the β pincer and sequence insertion βSI1^*63*^, which should affect formation of I_1E_ from RP_C_ and could also affect conversion of I_1M_ to I_1L_.

**Figure 7.**
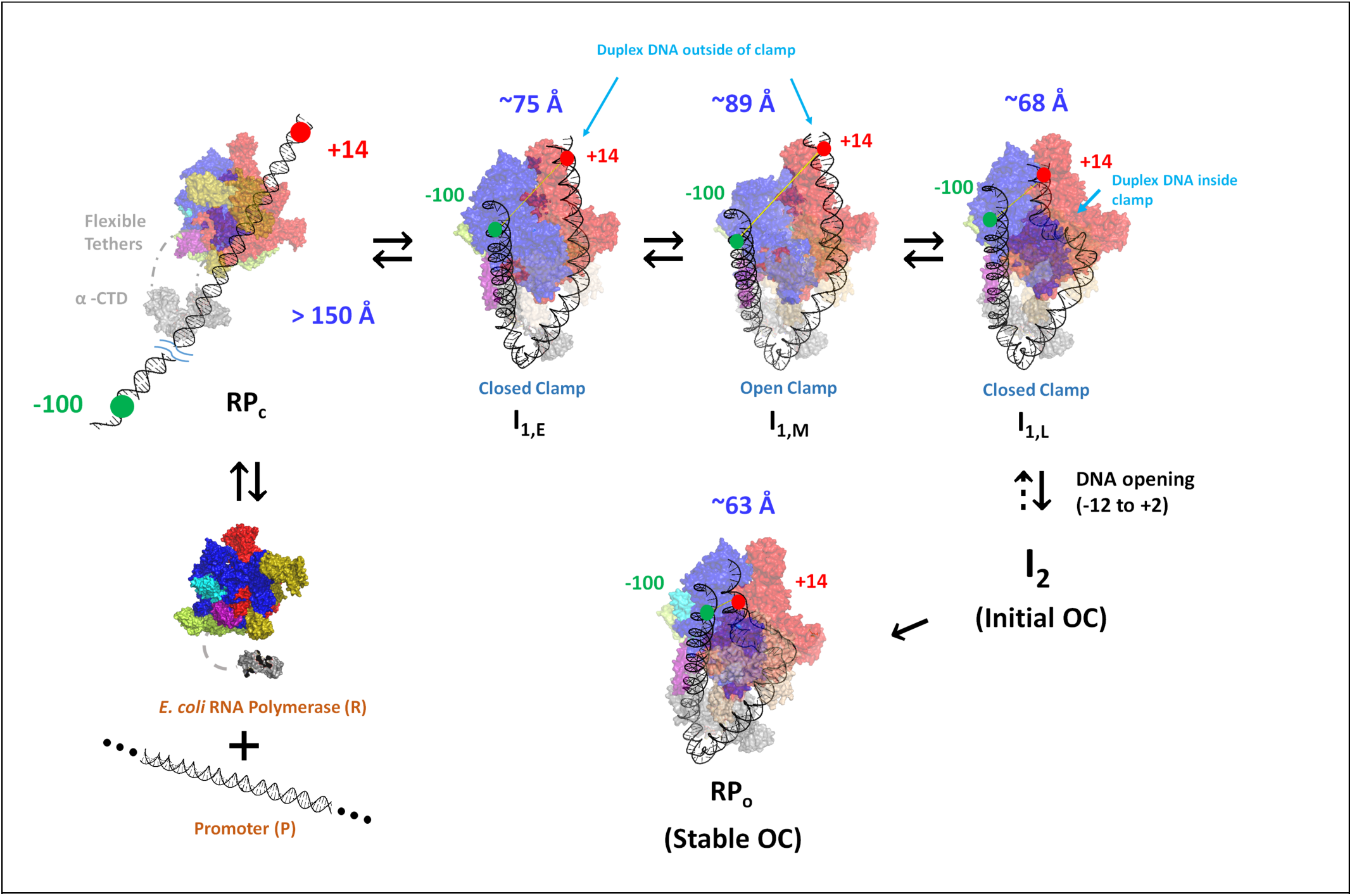
Illustrations of Bending and Wrapping of Promoter DNA in CC Intermediates and OC from FRET Distances and PIFE Contacts. Proposed mechanism of OC formation by RNAP determined by FRET and PIFE probes at −100 and +14 on promoter DNA. Unbound reactants (unbent promoter DNA, free RNAP) are at lower left. The absence of FRET and PIFE in the initial CC (RP_C_; upper left) shows that promoter DNA is not yet bent and wrapped on RNAP. Promoter DNA in subsequent CC intermediates (I_1E_, I_1M_, I_1L_) and in OC is highly bent and wrapped, making RNAP-DNA contacts at both −100 and +14. To explain the FRET distances and PIFE effects, we propose that concerted upstream bending/wrapping of upstream duplex DNA around the α subunits and onto the upper β’ clamp and bending of downstream duplex DNA onto the top of the β clamp in I_1E_ (top left-center) trigger clamp opening to form I_1M_ (top right-center). Clamp-opening triggers descent of the downstream duplex into the clamp and clamp-closing to form I_1L_ (top right), which opens the initiation bubble to form the initial unstable OC (I_2_). Stabilization of I_2_ by binding of RNAP mobile elements to the downstream duplex yields the stable OC (bottom right).

#### b) RNAP Clamp-Opening Moves Wrapped Upstream DNA Away from Downstream DNA to Form Low-FRET, High-PIFE I_1M_

The FRET efficiency of I_1M_ is less than that of I_1E_ (Table 2) while −100 and +14 PIFE signals of I_1M_ are similar to those of I_1E_ (Table 1). Analysis reveals that the conformational change that converts I_1E_ to I_1M_ increases the distance between the probes at −100 and +14 by ~14 Å (Table 2) without diminishing the PIFE contact of these positions with RNAP (Table 1). This increase in −100/+14 distance is readily explained if this step involves opening of the β’ clamp. Movement of the β’ clamp away from β in opening increases the distance between the tips of β and β’ pincers by ~14 Å^*64, 65*^. The time scale of clamp opening in the absence of promoter DNA (1 s)^*64, 65*^ is similar to that of the I_1E_ to I_1M_ conversion (Table 3). Fig. 7 therefore proposes that the far-upstream DNA is wrapped high on the outside of the upper portion of the β’ clamp, and that in clamp opening the −100 DNA moves ~14 Å away from +14 DNA on the top of the β subunit. This movement of the clamp presumably distorts the −10 region of the promoter DNA greatly.

#### c) The Downstream Duplex Is Bent into the Open Clamp, which Closes to Form the High-FRET, Most Advanced CC, I_1L_

The FRET efficiency of I_1L_ is greater than that of I_1E_ and much greater than that of I_1M_ (Table 2), indicating that the upstream DNA has moved much closer to the downstream DNA in I_1L_. We interpret this as a very large-scale set of conformation changes in which the downstream duplex is bent into the open clamp, followed by closing of the clamp (Fig 7). The −100/+14 distance is ~7 Å less than in I_1E_ and ~21 Å less than in I_1M_. A simple interpretation of these changes in distance is that bending the downstream duplex into the clamp brings +14 DNA ~7 Å closer to −100 DNA, and that closing the clamp brings −100 DNA ~14 Å closer to +14 DNA. The PIFE intensity at −100 is smaller for I_1L_ than for I_1M_ or I_1E_ (Table 1) indicating less strong contacts with both these DNA positions in I_1L_, while +14 PIFE intensity is similar for all three I1 intermediates.

#### d) Comparison with Other Mechanistic Proposals

A mechanism of open complex formation by E. coli σ^70^ RNAP was proposed based on fluorescence studies of interactions of designed short DNA oligomers with a −10 region or −35 and −10 regions (lacking upstream DNA), using RNAP drug complexes locked in closed and open clamp states. Probes of base flipping in the −10 element and of interactions of the +2 position in open complex formation were used in these studies^*46*^. Evidence was obtained for an early intermediate (called a recognition complex) in which closed-clamp RNAP is bound to somewhat bent, base-flipped but otherwise closed promoter DNA. This appears to correspond to the I_1E_ intermediate^*4*^ of Fig. 7 (without the upstream DNA) in which the RNAP is closed and the downstream DNA is sufficiently bent to contact the top of the β pincer. In the next proposed intermediate, called RP_I1_, the downstream duplex is more highly bent and the clamp of RNAP is open, similar to the I_1M_ intermediate characterized here. No evidence was obtained for the key late closed intermediate I_1L_. The initiation bubble in the next proposed intermediate (designated RP_I2_, based on a crystal structure of a fork-junction complex with open-clamp RNAP), appears similar to the initial, unstable OC previously identified by MnO_4_^−^ footprinting and kinetic analysis and called I_2_.^*52, 59, 64*^ Another possible I_2_ model is the open-clamp complex designated RP_ip_ observed by cryoEM with σ^54^ RNAP, stabilized by using a pre-melted heteroduplex DNA.^*51*^ Since we find the clamp closes in converting I_1M_ to I_1L_, the proposal that the clamp is open in I_2_ indicates that the clamp opens when the initiation bubble opens in converting I_1L_ to I_2_. If the strand backbones are held in the closed clamp in I_1L_, as seems likely, clamp opening may be connected to bubble opening in I_1L_ → I_2_.

Closed-promoter intermediates formed by yeast pol II RNAP have been characterized by cryoEM. In its downstream interactions, the I_1E_ intermediate proposed here (Fig. 7) is similar to the yeast pol II CC1 intermediate, in which the downstream duplex is bent and located above the closed clamp. In its downstream interactions, open-clamp I_1M_ (Fig. 7) corresponds to the subsequent pol II CC2 complex. In the next step for both enzymes, the downstream duplex is bent into the open clamp, but the clamp is closed on the downstream duplex in E. coli σ^70^ RNAP intermediate I_1L_ (Fig. 7) but the clamp is open in yeast pol II intermediate CC^dist^. Because the next step (DNA opening) for E. coli σ^70^ RNAP is rate-determining, no subsequent intermediate (like I_2_) is observable, but only the final OC. Presumably a similar situation exists for yeast pol II.

### Rate and Equilibrium Constants for Forming the Different Intermediates and their Use to Predict Time-Dependent Populations and the Free Energy vs. Progress Diagram

The time-evolution of populations of free promoter DNA (P), all four closed intermediates and OC for the reactant concentrations and conditions investigated here is shown in Fig. 8A. These predictions are based on the rate constants of Table 3, determined by fitting FRET and PIFE kinetic data to Mechanism 2.

**Figure 8.**
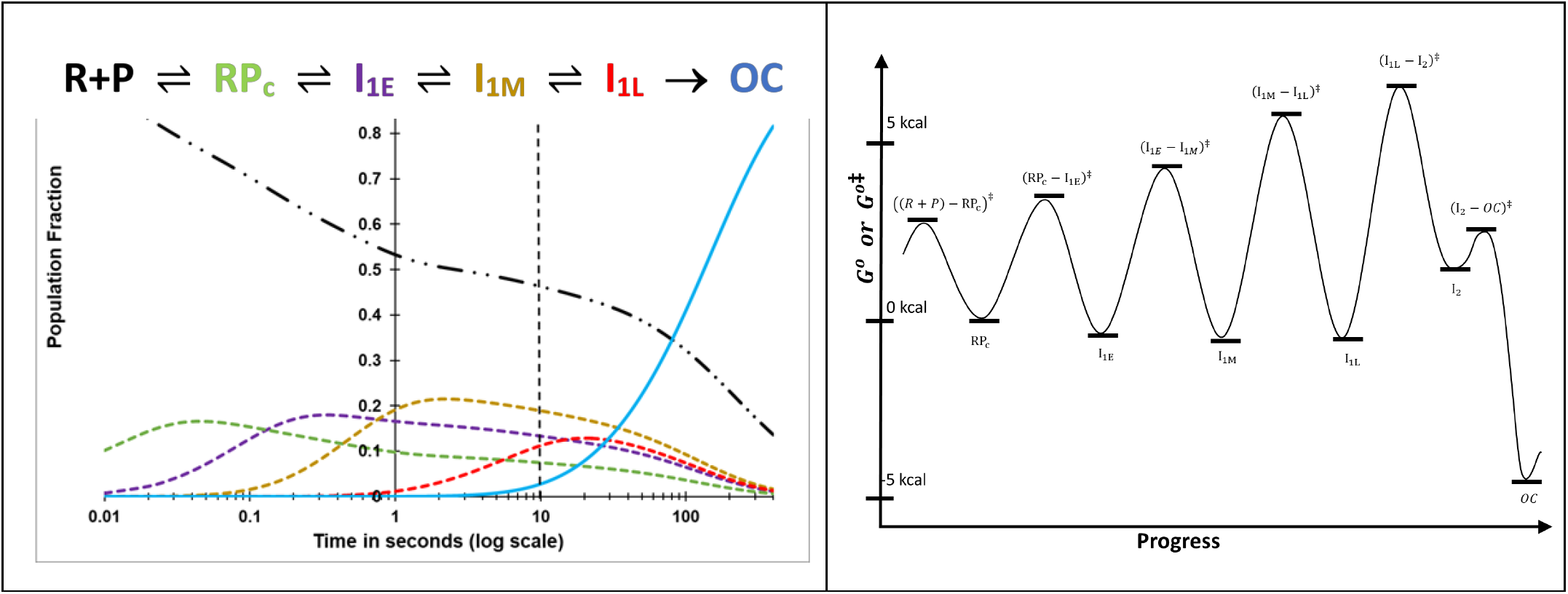
**A) Time Evolution of Populations of Unbound Promoter DNA, Intermediates in the {CC} Ensemble and the Stable λP_R_ Promoter OC.** Simulations of population fractions of reactant, intermediates and product vs time (0.01 s to 400 s) for 5-step Mechanism 2 at 50 nM final concentrations of RNAP and promoter DNA, 19 ° C, using rate constants (Table 3) from analysis of FRET and PIFE kinetic data. Free promoter DNA _. . _; closed complex intermediates RP_C_ _ _ _, I_1,E_ _ _ _, I_1,M_ _ _ _, I_1,L_ _ _ _,OC —. The dashed vertical line at 10 s marks the onset of OC formation from the equilibrium mixture of {CC} and free promoter DNA predicted from filter binding and MnO_4_^−^ kinetic data (Fig. 2B). **B) Standard Free Energy (G°) vs. Progress Diagram for OC Formation with FL λP**_**R**_ **Promoter DNA.** G° values for CC intermediates and the stable OC are obtained from the equilibrium constants of Table 3 (ΔG° = -RT lnK_i_) and are expressed relative to RP_C_, which is arbitrary assigned G° = 0 kcal. The G° value for I_2_ is obtained from K_5_ = k_5_/k_-5_, where k_-5_ = 1.2 s-1 (Fig. 5). Activation free energies G^*o*‡^ relative to the same reference are calculated from the relationship ΔG^*o*‡^ = - RTln(k/k _max_) where k_max_ is the (maximum) rate constant for the hypothetical situation ΔG^*o*‡^ = 0.^*4*^ For purposes of illustration, we choose k_max_ = 5 × 10^3^ s^-1^ for all steps.

### a) Free Promoter DNA (P)

Because the RP_C_ binding constant K_1_ is modest (~7 × 10^6^ M^-1^) and subsequent equilibrium constants for advancing RP_C_ to form the three closed I_1_ intermediates are also modest (in the range 1 – 2; Table 3), approximately half the population of promoter DNA remains unbound in the first 10 s of the reaction at the reactant concentrations simulated (50 nM). In these first 10 s, the {CC} forms as shown in Figs. 2B and 8A. After 10 s, formation of the very stable OC from the most advanced CC (I_1L_) results in additional conversion of P to RP_C_ and I_1_ species.

#### b) Initial Specific CC Intermediate (RP_C_)

Rate and equilibrium constants for RP_C_ formation (Table 3) are similar to published values for other promoters at similar conditions.^*49, 50*^ The second order rate constant k_1_ (~3.0 × 10^8^ M^-1^ s^-1^) for RP_C_ formation is about 5% of the diffusion-collision limit, most simply interpreted as that only about 5% of collisions of R with P result in RP_C_ formation. About 10% of promoter DNA is predicted to form RP_C_ in the first 10-20 ms of the reaction at the concentrations simulated (50 nM R, P). No FRET or PIFE signal is observed in this time range (Figs. 3, 4), and indeed none is expected for RP_C_ because footprinting indicates that the downstream promoter DNA is not bent and that the PIFE probe positions (−100, +14) are not in contact with RNAP. ^*8-12*^ The lifetime (1/k_-1_) of RP_C_ is relatively short (< 25 ms). RP_C_ increases to a maximum (~15% of total promoter DNA) at ~40 ms and remains the primary CC up to ~100 ms.

#### c) Wrapped CC Intermediate I_1E_

The rate constant for advancement of RP_C_ to I_1E_ (~13 s^-1^) is less than that for dissociation of RP_C_ (~ 44 s^-1^), allowing RP_C_ to equilibrate with R and P on the time scale of its conversion to I_1E_. I_1E_ is only marginally more stable than RP_C_ (K_2_ = 1.8 (Table 3); ΔG°_2_ = - 0.3 kcal/mol), showing that favorable contacts of these wrapped regions of promoter DNA with RNAP are just sufficiently favorable to overcome the costs of DNA-bending. Starting at 30 ms, the population of I_1E_ (Fig. 8A) increases to a broad maximum (about 20% of total promoter DNA) at 300 ms and then decreases gradually at longer times as more advanced CC and OC form.

#### d) Open-clamp Wrapped CC Intermediate I_1M_

The rate constant for advancement of I_1E_ to I_1M_ (~ 2.5 s^-1^) is less than that for reversal of I_1E_ to RP_C_ (~ 7 s^-1^), allowing I_1E_ to equilibrate with RP_C_ on the time scale of its conversion to I_1M_. I_1M_ is somewhat favored thermodynamically over I_1E_ (K_3_ = 1.5 (Table 3) and ΔG°_3_ = - 0.2 kcal/mol). The population of I_1M_ increases from ~100 ms to a broad maximum near 2 s (Fig. 8A) and then decreases gradually at longer times as more advanced CC and OC form. As in the previous step (RP_C_ → I_1E_), the cost of large conformational changes in conversion of I_1E_ to I_1M_ must be compensated by favorable interactions to result in a marginally favorable ΔG° for this step.

#### e) Closed-clamp Late CC Intermediate I_1L_

The rate constant for conversion of I_1M_ to I_1L_ (~0.1 s^-1^) is much smaller than that for reversal of I_1M_ to I_1E_ (~2 s^-1^), allowing equilibration of I_1M_ and I_1L_ on the time scale of conversion of I_1M_ to I_1L_. The population of I_1L_ increases from ~1 s to a broad maximum (~15% of total promoter DNA) at ~25 s and decreases gradually at longer times as I_1L_ slowly converts to OC in the DNA-opening step (Fig. 8A). The equilibrium constant K_4_ = 1.0 (Table 3) for conversion of I_1M_ to I_1L_ so these two intermediates have the same standard free energies (ΔG_4_° = 0).

#### f) Stable Wrapped OC

The FRET amplitude of the end-product 19 °C OC is somewhat larger than that of I_1L_, indicating that opening the bubble to form the initial unstable OC (I_2_) and subsequent formation of stabilizing downstream interactions in the conversion of I_2_ to the stable 19 °C OC reduce the distance between +14 and −100. Intrinsic PIFE signals at −100 and +14 in the stable OC are similar to those of I_1L_ and smaller than those of the earlier I_1_ intermediates (Table 1). Only overall changes in FRET and PIFE in converting I_1L_ to the stable OC are observed. While no information is obtained about the individual steps converting I_1L_ to I_2_ and then to the stable OC, previous quantitative kinetic studies of the steps of OC dissociation induced by KCl-upshift (as in Fig. 5) can be used to obtain this information.

#### g) Free Energy vs. Progress Diagram

Fig. 8B shows a standard free energy (G°) vs. progress diagram for Mechanism 2, based on the rate and equilibrium constants of Table 3. RP_C_ is taken as the point of reference and 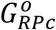 is set equal to zero. Activation free energies G^*o*‡^ relative to the same reference are calculated from the relationship ΔG^*o*‡^ = - RTln(k/k_max_) where k_max_ is the (maximum) rate constant for the hypothetical situation ΔG^*o*‡^ = 0.^*4*^ For purposes of illustration, we choose k_max_ = 5 × 10^3^ s^-1^ for all steps. The rapid-equilibrium condition for each step is indicated by the lower barrier for reversal of each intermediate, relative to going forward to the next intermediate. The highest G^*o*‡^ value is for the transition state for the rate-determining DNA opening step that converts I_1L_ to I_2_. Barriers surrounding I_2_ show the opposite pattern to those for previous intermediates, with a much lower barrier for forward conversion of I_2_ to the stable OC than for reversal of I_2_ to I_1L_.

Fig. 8B reinforces the finding (Table 3) that the steps starting with RP_C_ and forming the more advanced I_1_ intermediates are all only marginally favorable for this promoter and conditions, with equilibrium constants K_i_ between 1 and 2. Hence the net effect of bending and wrapping upstream and downstream promoter DNA to interact with the β’ and β subunits, respectively, only marginally favorable, and all I_1_ species are present at significant levels when the equilibrium distribution is established. This finding is consistent with HO footprinting results which showed that far-upstream contacts are substantially weaker than downstream contacts, and with the absence of a far-upstream DNase footprint, explicable because weak protein-DNA contacts can be displaced by binding of DNase.^*15*^

### How Far-Upstream DNA Facilitates Conversion of {CC} to OC: Insights from the Five-Step Mechanism

#### a) Upstream-DNA Greatly Increases the Isomerization Rate Constant k_isom_ at the λP_R_ Promoter with only a Modest Effect on K_CC_

The presence of upstream DNA (−40 to −100) in the FL λP_R_ promoter affects the kinetics of OC formation similarly to a strong, upstream-binding Type II transcription factor.^*16, 17*^ The isomerization rate constant of the FL λP_R_ promoter under the conditions investigated here is ~50 times larger than that of a λP_R_ promoter truncated upstream at position −47 (UT-47) ^*17*^ Similarly, k_isom_^FL^ for the lacUV5 promoter is 10-to 30-fold larger than k_isom_^UT^ for UT-42, UT-45, and UT-63.^*16*^ K_CC_ is either unaffected or increased by upstream truncation. ^*16, 17*^ These effects of the presence or absence of upstream DNA on k_isom_ of λP_R_ and lacUV5 promoters are as large as or larger than effects of promoter sequence changes or addition of upstream-binding factors.

From Eq. S19-20, upstream DNA could increase k_isom_ by increasing the fraction (f_I1L_) of the {CC} ensemble that is I_1l_ and/or by increasing the intrinsic DNA opening rate constant k_5_. *Several lines of evidence indicate that profound differences in f*_*I1L*_ *are the origin of the much greater k*_*isom*_^*FL*^ *as compared to k*_*isom*_^*UT 4*^. Real-time footprinting of {CC} ensembles during OC formation revealed that downstream contacts of RNAP with FL λP_R_ extend to +20 as compared to +2/+7 (partial protection) for λP_R_ UT-47^*15*^ and to −5 for RP_C_ at other promoters. To interpret these results we proposed that the UT-47 {CC} ensemble is composed primarily of RP_C_ and the subsequent CC intermediate (I_1E_), with relatively small concentrations of the more advanced CC (I_1L_ and probably also I_1M_) that are a major part of the FL {CC} ensemble^*4*^. We deduce that formation of these advanced CC from I_1E_ is favored by the presence of upstream DNA for the FL promoter and greatly disfavored for the UT promoter variants. Insights from Mechanism 2 and Fig. 7 help clarify why wrapping of FL upstream DNA is needed to advance the {CC} ensemble beyond I_1E_ and form a significant population of I_1L_, the only CC intermediate that can undergo the DNA opening step.

#### b) Predicted Large Differences in the Fraction of Advanced CC (f_I1L_) between FL and UT-47 λP_R_ Promoter DNA

Fractional populations of the different CC as a function of time in OC formation with full-length (FL) λP_R_ promoter DNA are shown in Fig. 8A. After equilibration (at > 10 s), the {CC} ensemble formed from FL λP_R_ promoter DNA is relatively advanced (33% I_1L_, 33% I_1M_) with smaller percentages of RP_C_ (12%) and I_1E_ (22%). This population distribution is consistent with real-time footprinting of the {CC} ensemble, which shows strong protection downstream to +20, characteristic of I_1L_ but not of I_1E_ or RP_C_^*15*^. By contrast, for UT-47 λP_R_, the analysis given in SI predicts that I_1L_ and perhaps also I_1M_ are greatly destabilized (Fig. 9). The predicted population distribution in the equilibrated {CC} ensemble for UT-47 is approximately 60% I_1E_, 33% RP_C_, 6% I_1M_ and only 0.6 % I_1L_. This small population of I_1L_ in the equilibratedUT-47 {CC} ensemble is the likely origin of its small isomerization rate constant.

**Figure 9.**
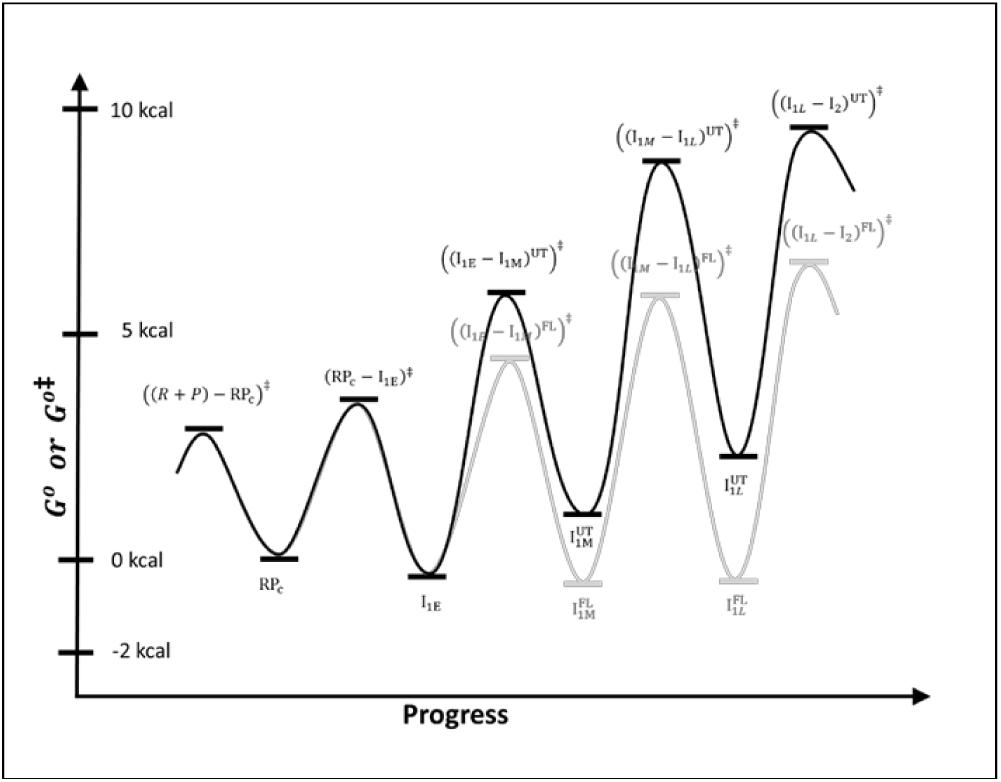
Proposed Standard Free Energies (G°) vs. Progress Diagram for OC Formation with UT-47 λP_R_ Promoter DNA and Comparison with FL λP_R_ Promoter DNA. Standard free energies (G°) for CC intermediates formed by UT-47 λP_R_ promoter DNA are obtained from equilibrium constants as described in SI and are expressed relative to RP_C_, which is arbitrary assigned G° = 0 kcal. Effects of truncation on the steps converting I_1E_ to I_1L_ are assumed to be entirely on the forward rate constant of these steps. Activation free energies G^*o*‡^ relative to the same RP_C_ reference are calculated from the relationship ΔG^*o*‡^ = - RTln(k/k_max_) where k_max_, the (maximum) rate constant for the hypothetical situation ΔG^*o*‡^ = 0, is arbitrarily assigned the value k_max_ = 5 × 10^3^ s^-1^ for all steps. Diagram from Fig. 8B for the full-length (FL) promoter is shown for comparison.

#### c) Proposed Structural Basis of these Very Different {CC} Population Distributions

For FL λP_R_ the kinetic data and above analysis show that one or both steps of conversion of I_1E_ to I_1L_ is/are greatly favored by the interactions of bent-wrapped upstream DNA with RNAP, as previously proposed^*4*^. If conversion of I_1E_ to I_M_ is the step that is much more favorable for FL λP_R_ than for UT-47, then bending and wrapping of upstream DNA on RNAP favors clamp opening. This might involve compaction or folding of the α-CTD tethers to bend and wrap the upstream DNA in I_1E_, causing the assembly of UP element DNA, α-CTD, and compacted tethers to interact with the body of RNAP. A structural analogy would be the lac repression complex, in which the flexible tethers connecting the DNA binding domains (DBD) to the core repressor fold into helices, interact with operator DNA, and cause the DNA-DBD assembly to interact with the repressor core^*66-69*^.

If conversion of I_1M_ to I_1L_ is the step that is much more favorable for FL λP_R_ than for UT-47, then upstream wrapping favors the entry and descent of the downstream duplex into the open clamp. A molecular explanation for this, proposed previously^*3, 15*^, is that upstream wrapping could allow far-upstream DNA to contact downstream mobile elements (DME) of RNAP and move them away from the downstream clamp/cleft. This proposal and the interaction of far upstream DNA with DME would explain the significant −100 PIFE signals of I_1E_ and I_1M_ (Table 1), and the reduction in −100 PIFE in I_1L_ and OC when the DME move to interact with the downstream duplex in the stable OC ^*4*^.

We think it likely that both these steps (I_1E_ → I_1M_ and I_1M_ → I_1L_) are affected by contacts of the upstream-wrapped DNA with the outside of the clamp and with the DME at the downstream end of the clamp (Fig. 7), and therefore that both conversions are facilitated by upstream wrapping. An example of this is shown in the comparison of free energy vs. progress diagrams for UT-47 and FL λP_R_ in Fig. 9 (details provided in SI). Fig. 9 predicts that the much smaller k_isom_ for UT-47 results from the relative instability of late CC intermediates 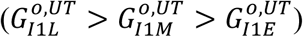, in contrast to the situation for FL λP_R_ where late CC intermediates are more stable than earlier ones 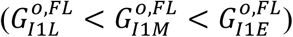.

## Conclusions

The FRET and PIFE fluorescence kinetics studies reported here reveal for the first time when upstream and downstream promoter DNA are bent and wrapped on RNAP in the mechanism of OC formation, as well as why upstream DNA (−40 to −100) is necessary for efficient isomerization of the {CC} ensemble to OC. Information in the promoter sequence (presumably primarily the UP-element, −35 and −10 regions), read by interactions with the α-CTD and σ^70^ regions 4 and 2 in the initial CC (RP_C_), directs concerted bending and wrapping of both upstream and downstream DNA in conversion of RP_C_ to the more advanced CC, designated I_1E_, in which the distance between −100 and +14 positions on promoter DNA is reduced from >300 A before binding of RNAP to ~75 A. Interactions between promoter DNA and RNAP in I_1E_, which we propose are primarily between the upstream-wrapped DNA and the hinge of the clamp, result in opening of the clamp (on a ~1 s timescale for the conditions investigated) to form the intermediate I_1M_. Clamp opening moves the far-upstream DNA, which we propose is wrapped high on the back of the β’ clamp, ~14 A further away from the downstream duplex, which we propose is bound on the top of the β clamp in I_1E_ and I_1M_. Subsequently, on a ~10 s timescale, the downstream duplex is bent into the clamp, which closes to reduce the distance between −100 and +14 positions to ~68 A in I_1L_, the most advanced CC. Opening of the initiation bubble by strand binding free energy appears to occur in a single kinetic step at this promoter to form the initial unstable OC (I_2_), which is rapidly stabilized by downstream interactions with DME. Each step of forming and advancing the {CC} ensemble rapidly equilibrates on the time scale of the next step. The DNA opening step is rate-determining for OC formation because its forward rate constant (k_5_ = 0.04 s^-1^) is smaller than those of previous steps and because it is effectively irreversible since stabilization of I_2_ by downstream interactions is more rapid than DNA-closing for the promoter and conditions investigated (Fig. 8B).

## Acknowledgments

We thank previous undergraduates Emily Lingemann, Miranda Mecha, Yurun Zhang, Priya Chittur, Wen Fu, Rahul Vivek, and Hertina Kan for their contributions to this research. We thank Prof. J. Weisshaar for his comments on the manuscript, and gratefully acknowledge support of NIH grant GM R35-118100.

## Accession Codes

P0A7Z4, subunit *α*; P0A8V2, subunit *β*; P0A8T7, subunit *β’*; P0A800, subunit *ω*; P00579 subunit *σ*.

### Contents of SI

The SI text describes the methods used in the following analyses. 1) the normalization of fluorescence kinetic data; 2) fitting methods, including the analysis of FRET and PIFE kinetics as a sum of four exponentials and fitting these data to five-step Mechanism 2; 3) consistency checks on the rate constants of Mechanism 2; 4) conversion of fluorescence fitting amplitudes to relative fluorescence intensities; 5) estimation of distance between upstream (−100) and downstream (+14) DNA in wrapped RNAP-promoter complexes from FRET amplitudes; 6) interpretation of rate and equilibrium constants of the classic two-step Mechanism 1 (k_isom_, K_CC_, k_a_) using Mechanism 2; and 7) calculations for free energy vs. progress diagram and {CC} population distribution for formation at UT-47 λPR. SI figures include: 1) the MnO_4_^−^ footprinting data of Fig. 1 plotted on a logarithmic time scale; 2) comparison of the FRET and PIFE data of Figs. 3-4 with the prediction based on the average rate constants and signal amplitudes of Tables 1 and 3. SI Tables include: 1) promoter and 2) primer sequences used in PCR synthesis of the different dye-labeled promoters investigated here; 3) comparison of rate constants (1/*τ*_*i,obs*_) from the four-exponential fit with predictions from the rate constants of Mechanism 2; 4) individual rate constants and signal amplitudes obtained from analysis of all FRET and PIFE experiments; and 5) average FRET acceptor and PIFE fitting amplitudes of CC intermediates and the stable OC.

## Supplemental Information

### Normalization of Fluorescence Kinetic Data

For each FRET and PIFE OC formation experiment, a time-averaged initial (t ≈ 10 ms) minimum fluorescence (F_min_) and a maximum fluorescence (F_max_) in the interval 10 ms to 400s were obtained for each shot and used to calculate a normalized fluorescence (F_norm_) as a function of time using Eq. S1.

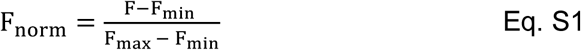

Visual inspection of the set of 10-15 normalized shots in each series was used to eliminate any systematic outliers. Then these normalized data were averaged for subsequent fitting.

In previous FRET-detected equilibrium titrations of these Cy3Cy5 labeled promoter DNAs (at 50 nM) with RNAP, evidence for a competitive nonpromoter binding mode was obtained at RNAP:promoter DNA ratios greater than 1:1.^1^ To avoid this, kinetics experiments were performed at concentration ratios of active RNAP to promoter DNA in the range 0.5:1 to 1:1.

### Fitting Methods

#### Analysis of FRET and PIFE Kinetics as Sums of Exponentials

FRET and PIFE OC formation kinetic data (each an average of a series of ~10 normalized shots) collected over the time range 10 ms – 400 s were fit with Berkeley Madonna^2^ to a sum of four exponentials and a linear correction for long-term drift^3^ (Eq. S2):

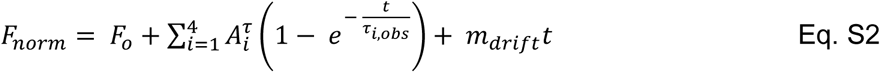

where the *τ*_*i,obs*_ (i = 1 − 4) are time constants and 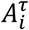 are amplitudes. F_o_ is the extrapolated fluorescence at t = 0 (negative because of the normalization at 10 ms; Eq. S1) and *m*_*drift*_ is the slope of the drift correction, which generally was small and varied from experiment to experiment. We found that three exponentials in Eq. S2 were insufficient while five were too many, supporting the use of Mechanism 2 with three detectable intermediates, differing in FRET and PIFE from the final open complex. Rate constants 1/*τ*_*i,obs*_ were obtained from the fits to Eq S2. Average values of 1/*τ*_*i,obs*_ calculated from all FRET and PIFE experiments reported here and listed in Table S3, span four orders of magnitude, in decade increments from ~ 20 s^-1^ to ~ 0.02 s^-1^. No significant differences in 1/*τ* _,*obs*_ values were detected for the various types of FRET and PIFE experiments.

Because filter binding and MnO_4_^−^ kinetics of OC formation are single-exponential (see for example Figs. 1 and S1), the closed complex ensemble {CC} in Mechanism 1 must rapidly equilibrate with P and R on the time scale of isomerization of {CC} to OC, and therefore each step of forming and advancing the CC ensemble in Mechanism 2 must equilibrate rapidly on the time scale of the next step. For this reason values of 1/*τ*_*i,obs*_ (i ≤ 3) are interpretable as decay-to-equilibrium rate constants^4^ for the steps in Mechanism 2 detected by FRET and PIFE that form the three I_1_ intermediates.

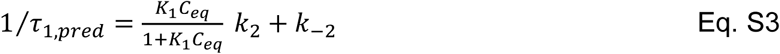

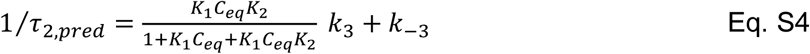

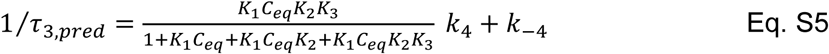

where C_eq_ = ([P]_eq_ + [R]_eq_).

The smallest rate constant 1/*τ*_4,*obs*_ is interpreted as the rate constant for the subsequent, effectively-irreversible DNA opening step that converts the most advanced I_1_ intermediate (I_1L_) to I_2_.

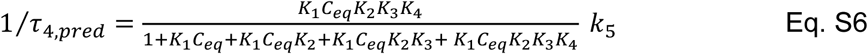

In fitting to Mechanism 2, initial order-of-magnitude estimates of rate constants were obtained from the corresponding 1/*τ*_*i,obs*_ values (i ≤ 3) assuming k_i_ ~ k_-i_ ~ 1/(2*τ*_*i*−1,*obs*_) and k_5_ ~ 1/*τ*_4,*obs*_. Semi-quantitative comparisons of values of 1/*τ*_*i,obs*_ obtained from fitting FRET and PIFE data to Eq S2 with values obtained from equilibrium and rate constants from fits of these data to Mechanism 2 are given in Table S3. In the time range from ~1 s to ~10 s which is of most interest for these comparisons, C_eq_ ≈ [P]_tot_ based on the simulations in Figs. 2B and 7 which show that [P] = [R] ≈ 0.5[P]_tot_ in this time range.

#### Fitting to Mechanism 2

The normalized fluorescence *F*_*norm*_ is interpreted as a sum of contributions from the I1 intermediates and OC in Eq. S7.

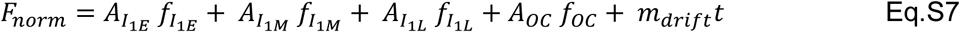

where 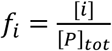 is the time-dependent population fraction of species i (i = I_1E_, I_1M_, I_1L_, or OC) with intrinsic signal amplitude *A*_*i*_ and *m*_*drift*_ is the slope of the linear drift correction. Justification for this interpretation of *F*_*norm*_ is provided in the next section (Eqs. S8-14).

Values of F_norm_ vs. time from the high signal-to-noise (high S/N) Cy5 FRET acceptor experiments in Fig. 3, which exhibit little if any drift when fit to Eq. S2, were fit interactively to Mechanism 2 using Kintek Explorer^5,6^ with the constraints described below to determine the nine rate constants k_i_ and the FRET acceptor amplitudes *A*_*i*_ of the three I_1_ intermediates and the stable OC. Analyses in which a possible FRET contribution from RPC was included revealed that 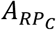 is small and not significantly different from zero, consistent with it being the initial, not-yet-wrapped closed complex.

As initial estimates of FRET amplitudes to use in fitting to Mechanism 2, we assumed a monotonic increase in FRET and arbitrarily set 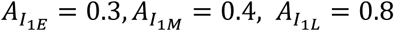. The choice *A*_*oc*_ = 1.2 is based on the normalization *F*_*norm*_ = 1.0 (Eq. S1) at t = 400 s, where from the simulation in Fig 2B the population is approximately 80% OC. Initial estimates of rate constants for steps 2-5 of Mechanism 2, in which RP_C_ is converted to OC via the series of I_1_ intermediates, were obtained from the corresponding 1/*τ*_*i,obs*_ values as described above. These initial estimates provide rapid equilibration of each step on the time scale of the next. Initial estimates of rate constants for the first step were k_1_ ~ 6 × 10^8^ M^-1^s^-1^ and k-1 ~ 60 s^-1^, consistent with rapid equilibrium of this step on the time scale of the next, and consistent with an equilibrium constant K_1_ = K_RPc_ of ~10^7^ M^-1^ as obtained from K_CC_ (Fig. 2B) and initial estimates of the subsequent equilibrium constants (K_i_ ≈1-2 for 2 ≤ i ≤ 4)) using Eq.S21-S22.

Reasonable fits to *F*_*norm*_ were obtained with fitted values of rate constants that differed from the initial estimates while maintaining a hierarchy so that each step equilibrated rapidly on the time scale of the next. Relative amplitudes of intermediates I_1E_ and I_1M_ inverted in these fits from the initial estimates so 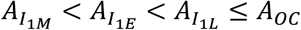. However these initial fits failed to satisfy various constraints. In particular, the fraction of P near the end of the {CC} phase (t ~ 1 s) and/or the fraction of OC at late in OC formation (400 s) did not agree with the simulation in Fig 2B. Also, fitted rate constants for one or more intermediate steps differed significantly from expectations based on 1/*τ*_*i,obs*_ values.

To refine the fit, pairs of rate constants k_-i_, k_i+1_ were linked by constraining each k_-i_ to be at least two-to-three times greater than k_i+1_ in order to have each step equilibrate on the time scale of the next. Then in interactive fitting k_1_ and subsequent forward rate constants were varied to obtain a fraction of free promoter DNA (i.e. [P]/[P]_total_) at 1 s of approximately 0.5 and a fraction of promoter DNA present as OC (i.e. [OC]/[P]_total_) at 400 s of approximately 0.8, as expected from the simulation in Fig. 2B. Optimization of these fits yields the rate constants and species amplitudes in Table S4, which provide high-quality fits to the data of Fig. 3 as shown in Fig. 6. In these optimized fits, the rank order of FRET amplitudes is 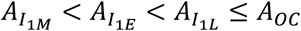 for both Cy5-100 and Cy5+14 FRET experiments. These results differ from our initial assumption in fitting these data of a monotonic increase in FRET amplitude in the steps of OC formation.

To analyze the remaining FRET and PIFE data sets, many of which exhibited lower S/N ratios and/or larger drift than in the experiments analyzed above, rate constants were allowed to vary by up to ±10% from those obtained in the above analysis, and amplitudes of intermediates and OC were allowed to vary without any restriction in Berkeley Madonna^2^ fitting. Global fitting of all FRET or PIFE kinetic data obtained for the same probe position(s) was not feasible because of the lower S/N ratio of many experiments, differences in the normalized signal at longer times (t> 50 s) in PIFE experiments, even after drift-correction, and because of differences in time points of data acquisition. High-quality fits over the entire time-range were obtained for more than 90% of the 61 data sets. Average amplitudes and uncertainties for each species detected are reported in Table S5. More than 75% of individual FRET acceptor experiments exhibit the same rank order of FRET amplitudes 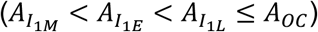 and the others show only a single deviation from this order. Experimental FRET and PIFE kinetic curves from Figs. 3 and 4, corrected for linear long-time drift where necessary, are compared with those predicted from average rate constants (Table 3) and amplitudes (Table S5) in Fig S2.

#### Consistency Checks on the Rate Constants of the 5-Step Mechanism

Rate constants and equilibrium constants for the steps in Mechanism 2 (Table 3) pass several consistency tests.

a. Use of these rate and equilibrium constants in decay-to-equilibrium analyses of the reversible steps of Mechanism 2 (Eqs. S3-S7), predict the rate constants (1/*τ*_*i,obs*_ values) obtained from empirical 4-exponential fits to the FRET and PIFE kinetic data (Table S3).
b. These rate and equilibrium constants are quantitatively consistent with the filter binding and MnO_4_-results of Fig. 2. Eqs. S21-22 predict the composite {CC} binding constant K_cc_ (Fig. 2A) of Mechanism 1 from the equilibrium constants K_RPc_, K_2_, K_3_ and K_4_ of Mechanism 2 (in Table 3). Eqs. S19-20 predict the isomerization rate constant k_isom_ of Mechanism 1 from the DNA opening rate constant k_5_ and equilibrium constants _K2_, K_3_ and K_4_ of Mechanism 2 (in Table 3). Predicted and experimental values of K_CC_ and k_isom_ agree within the uncertainty, as shown in Table 3 footnotes.
c. Values of equilibrium constants for the steps converting RP_C_ to I_1L_ in the range 1 – 2 are consistent with real-time hydroxyl radical footprint of the equilibrated {CC} ensemble under similar conditions to those investigated here.^7^ These exhibit a downstream boundary at +20, like that of the stable OC, indicating a dominant population of I_1_ (i.e. post-RP_C_) intermediates and hence values of K_i_ ≥ 1 for at least some of these steps (1< i < 4). Also, downstream protection for this {CC} is significantly weaker than for OC, and the far-upstream footprint is also of modest intensity, indicating that these contacts are not highly stable like those of the −10 and −35 elements and therefore that values of K_i_ for these steps are not much greater than 1. Rapid equilibrium also requires that values of Ki for these steps cannot be much greater than 1.
d. 5-step Mechanism 2 allows a quantitative interpretation of the large effect of upstream truncation (e.g. UT-47) and consequent loss of far-upstream wrapping on k_isom_ and the small effect of upstream truncation on K_cc_^8^, using Table 3 rate constants for steps that are proposed to be unaffected by truncation, as discussed in a subsequent section.

#### Conversion of Fluorescence Fitting Amplitudes to Relative Fluorescence Intensities

Here we obtain Eq. S7, used to fit normalized FRET acceptor and PIFE kinetic data, and interpret species amplitudes *A*_*i*_ obtained from these fits in terms of their fluorescence intensities *a*_*i*_. The unnormalized fluorescence signal F, after correction for drift (if any) at long times, is the sum

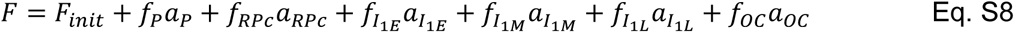

where ai is the fluorescence intensity of species i and 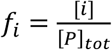 is its concentration fraction. In Eq. S8, *i* represents P, RP_C_, I_1E_, I_1M_, I_1L_, or OC and ∑_*i*_ *f*_*i*_ = 1.

For the present FRET and PIFE applications,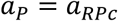 and 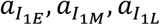 and *a*_*oc*_ all are larger than *a*_*P*_ and *a*_*RPc*_. We define intensity differences Δ*a*_*i*_ between each species i and free promoter P:

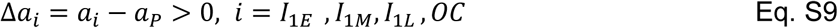

Because 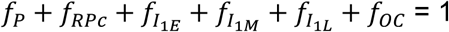, therefore

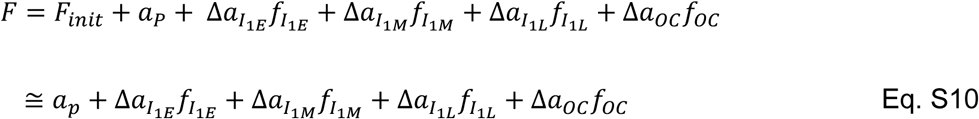

because the background fluorescence *F*_*init*_ is small in comparison to the fluorescence *a*_*P*_ of P (*Fi*_*init*_ ≪ *a*_*P*_).

The minimum fluorescence F_min_ (Eq. S1) is the fluorescence at t_0_, the initial time (~10 ms) of data collection. Because no significant amount of any fluorescence-detected (i.e. I_1_) intermediate is present at time t_0_ (see Fig. 7), therefore 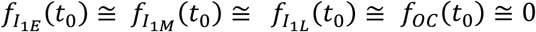 and from Eq. S10:

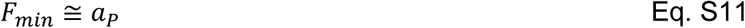

The maximum fluorescence F_max_ (Eq. S1) occurs at time t_max_.

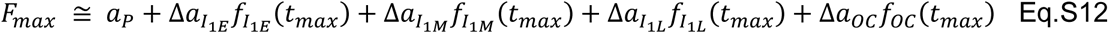

Therefore the normalized fluorescence *F*_*norm*_ is given by

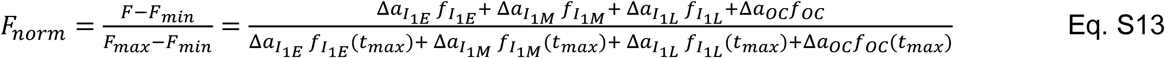

which is the same as Eq. S7 with

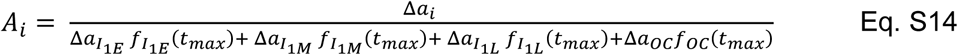

where *i* represents. *I*_1*E*_,*I*_1*M*_,*I*_1*L*_ or *OC*

From Eqs. S11-12,

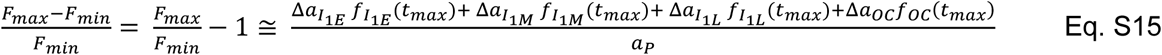

and from Eqs. S14-15

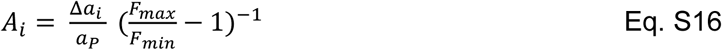

The average value designated 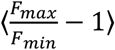 is obtained by first averaging factors 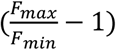 for the series of shots in each experiment and then averaging over all PIFE experiments of a given type. These 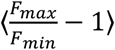 values and average fitting amplitudes ⟨*A*_*i*_⟩ are reported in Table S5, together with uncertainties in these quantities. Average PIFE intensities 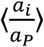 relative to promoter DNA are obtained from these quantities by Eqs. S17 (the average of Eq. S16) and reported in Table 1.

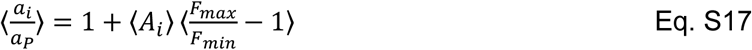

#### Estimation of Distance between Upstream (−100) and Downstream (+14) DNA in Wrapped RNAP-promoter Complexes from FRET Amplitudes

Cy5+14 and Cy5-100 FRET fitting amplitudes (Table S5) were normalized by the OC fitting amplitude to obtain the values listed in Table 1. These are averaged and converted to FRET efficiency values of intermediates I_1E_, I_1M_ and I_1L_ in Table 2, using the published OC FRET efficiency (0.32).^1^ These FRET efficiencies are converted to distances using Eq. S18:

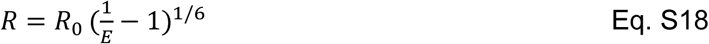

with the Forster radius previously estimated for this DNA at 19 °C (R_0_ = 56 ± 1 A). Illustrations of intermediates with these FRET distances are shown in Fig. 8.

#### Interpretation of Rate and Equilibrium Constants of the Classic Two-Step Mechanism 1 (k_isom_, K_CC_, k_a_) Using Mechanism 2

Previously, effects of upstream DNA truncation, promoter sequence and upstream-binding factors on the kinetics of OC formation were interpreted using 2-step Mechanism 1 as effects on the isomerization rate constant k_isom_ and on the composite equilibrium constant for formation of the {CC} ensemble, K_CC_. Interpretation of these effects using relationships between these parameters of Mechanism 1 and the rate and equilibrium constants (Table 3) of 5-step Mechanism 2 provides new insight into how promoter sequence, upstream truncation and factors regulate the rate of OC formation and transcription initiation.

Because each step of {CC} ensemble formation equilibrates rapidly on the time scale of the next, k_isom_ of Mechanism 1 is related to k_5_, the rate constant of the DNA-opening step in Mechanism 2, by

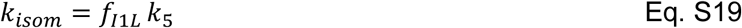

where f_I1L_ is the fraction of the equilibrated {CC} ensemble that is I_1L_:

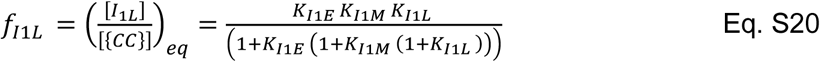

where

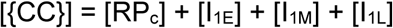

The {CC} ensemble is equilibrated at t ≥ 10 s (the onset of OC formation) for the conditions investigated here (Fig 7).

For this rapid equilibrium situation, K_CC_ of Mechanism 1 is related to the equilibrium constant for forming the initial closed complex, K_RPc_, by

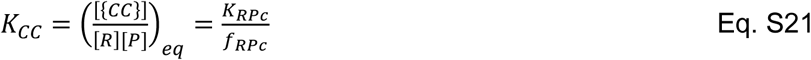

where f_RPc_ is the fraction of the equilibrated {CC} that is RP_C_:

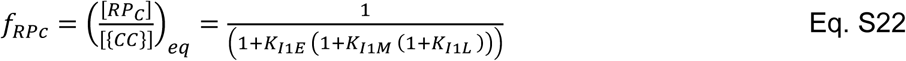

Eqs. S19-22 show that k_isom_ < k_5_ and K_CC_ > K_RPc_. The overall second order rate constant for OC formation (k_a_) is interpreted using these two mechanisms as

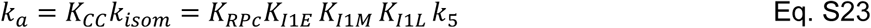

In the equilibrated {CC} ensemble, population fractions of I_1E_ and I_1M_ intermediates are

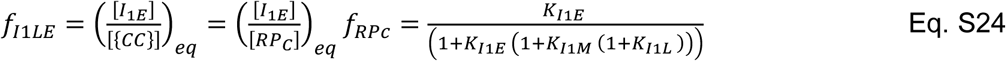

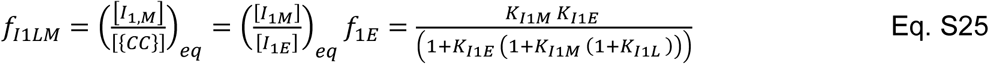

Effects of promoter sequence, upstream DNA length and transcription factors on the parameters of Mechanism 1 (k_isom_, K_CC_, and k_a_) can arise from effects of these variables on the distribution of complexes in {CC} (i.e. on f_RPc_ and f_I1L_) as well as from effects of these variables on K_RPc_ and k_5_. Based on the limited evidence available to date, and by analogy with regulation of enzyme catalysis, we proposed previously that the 100-fold or greater range of k_isom_ for different upstream promoter truncations, different promoter sequences, and/or addition of transcription factors results primarily from an equally wide range in the fraction f_I1L_ of the equilibrated {CC} ensemble that is I_1L_, the intermediate that is poised for DNA opening, and not primarily from changes in the DNA opening rate constant k_5_ with these variables.^9^ An example is provided in the Discussion. This proposal is supported by the finding that, for λP_R_ and T7A1 promoters and wild-type and variant RNAP, k_-5_, the DNA closing rate, is insensitive to promoter identity and to salt or solute concentration, and primarily a function of temperature^10-12^

A fundamental analogy exists between Mechanism 2 for open complex formation in the RNAP clamp and mechanisms of enzyme action. Regulation of the velocity of product formation in noncooperative and cooperative enzyme-catalyzed reactions is often achieved using inhibitors or activators that primarily affect the equilibrium constant(s) for reversible binding of the substrate(s), and not the catalytic rate constant. Likewise, effects of promoter sequence and transcription factors on the rate of OC formation appear to be largely on equilibrium constants for the steps that form RPc and convert it to the more advanced members of the CC ensemble (I_1E_, I_1M_, I_1L_). These reversible steps are analogous to reversible substrate binding steps in enzyme kinetic mechanisms, and in-cleft DNA opening-closing is the analog of the catalytic step.

#### Calculations for Free Energy vs. Progress Diagram and {CC} Population Distribution for Formation at UT-47 λP_R_

Assuming the same DNA-opening rate constant (k_5_) for UT-47 and FL λP_R_ at 19 °C, then from the experimental k_isom_^UT^ it follows from Eq. S19 that f_I1T_^UT^ = 0.006. Hence, less than 1% of the equilibrated UT-47 {CC} ensemble is I_1L_ as compared to 33% for FL, explaining the 60-fold difference in isomerization rate constants. The remaining 99% of the UT-47 {CC} ensemble is a mixture of RP_C_, I_1E_, and possibly I_1M_. The large reduction in k_isom_ for UT-47 indicates a large reduction in K_I1L_ and/or K_I1M_.

For the progress diagram in Fig. 9, we assume that K_I1E_^UT^ = K_I1E_^FL^ = 1.8 and that K UT = K_I1M_^UT^ << 1. In this case, we calculate K_I1M_^UT^ = K_I1L_^UT^ = 0.098 as compared to K_I1M_^FL^ = 1.5 and K_I1L_^FL^ = 1.0 for FL (Table 3). From these equilibrium constants, the fully-formed UT-47 {CC} ensemble is predicted to be 33% RP_C_, 60% I_1E_, 6% I1M, and only 0.6% I_1L_, consistent with the {CC} ensemble footprinting result for UT-47. For comparison, the {CC} ensemble for FL λP_R_ (Fig. 8A) is 12% RP_C_, 22% I_1E_, 33% I_1M_ and 33% I_1L_.

## SI Figures

**Figure S1:**
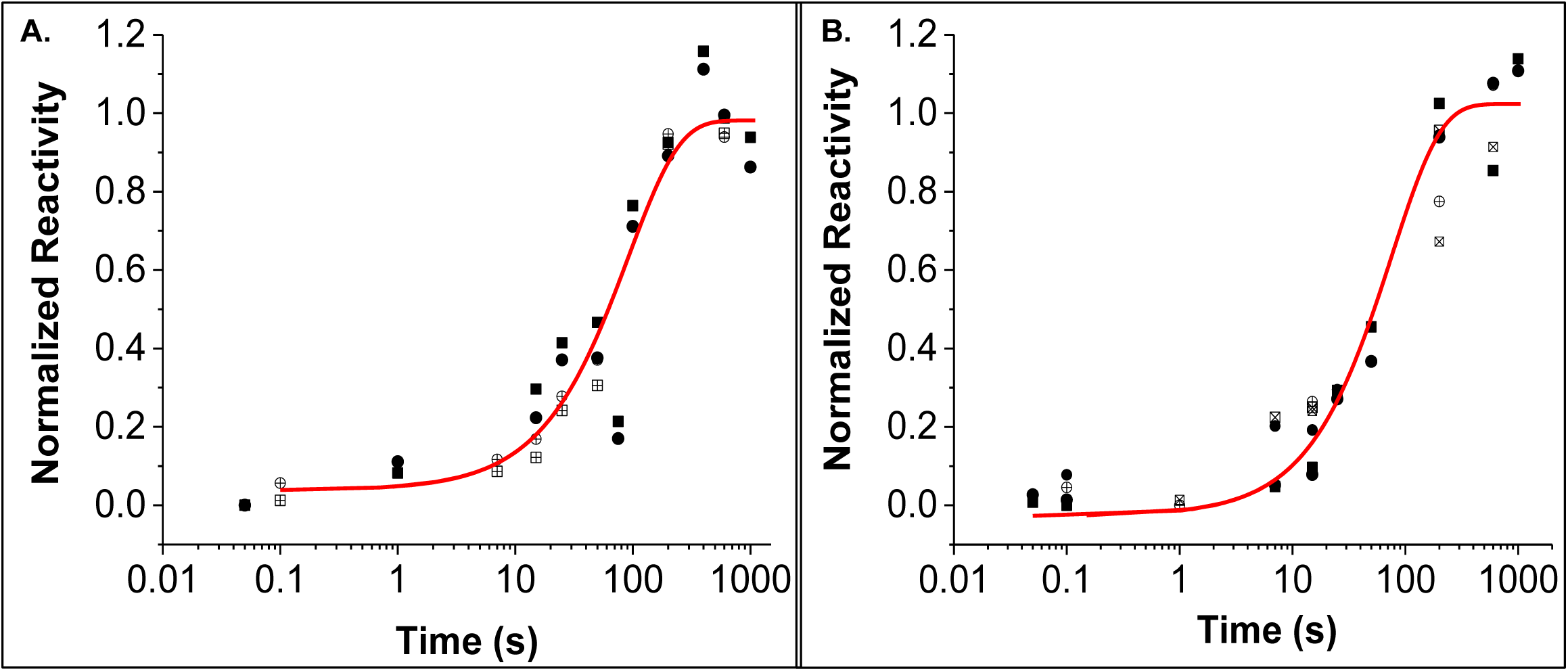
MnO_4_^−^-Detected Kinetics of OC Formation on a Log Time Scale. Results in Fig. 1 of the text for the kinetics of OC formation monitored by fast permanganate footprinting are replotted on a log time scale to ascertain if the entire time course is described by single-exponential kinetics. Symbols for different thymines and fitted curves are the same as in Fig. 1 and correspond to rate constants k_obs_ = 0.011 ± 0.002 s^-1^ (non-template strand, Panel A) and k_obs_ = 0.010 ± 0.002 s^-1^ (template strand, Panel B). These comparisons reveal that all reactive thymines on each strand exhibit very similar single-exponential kinetics of OC formation.

**Figure S2:**
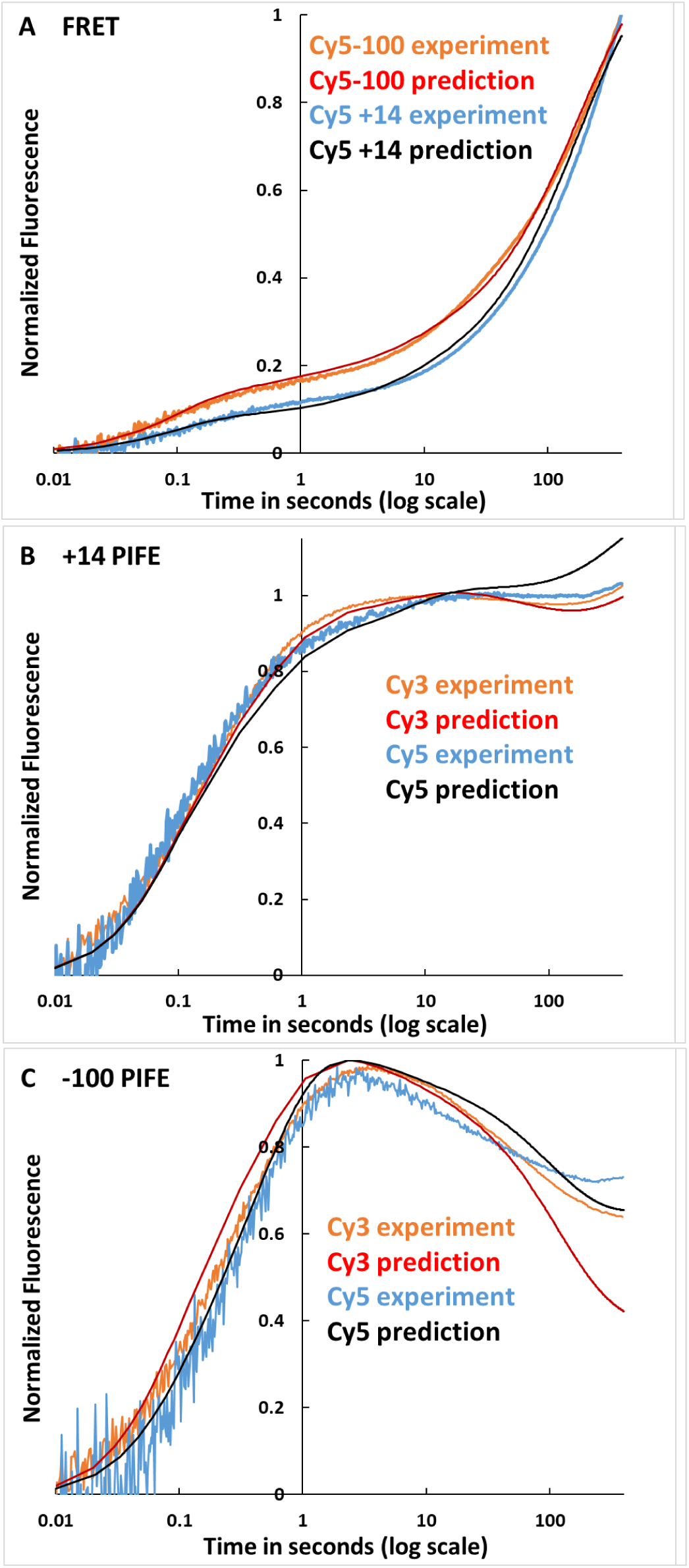
Comparison of Fitted (Predicted) and Observed FRET and PIFE Kinetics of OC Formation. Comparison of FRET and PIFE kinetic data (as in Figs. 3,4, but after a linear drift correction) with predictions using average rate constants (Table 3) and amplitudes (Table S5). **Panel A**: **FRET** for Cy5 −100 —; Cy5 +14 —. **Panel B**: **PIFE** for Cy3 +14 —; Cy5 +14 —. **Panel C: PIFE** for Cy3 −100 —; Cy5 −100 —. Elimination of signal drift from the +14 PIFE data (Figs. 4, 6) reveals a small increase in +14 PIFE at t >50 s. A similar but smaller effect is observed for −100 PIFE. The late +14 PIFE increase results from the conversion of remaining free P and RP_C_ (no PIFE intensity) to OC (modest PIFE intensity), which more than counterbalances the effect of converting higher-PIFE I_1_ intermediates to OC.

**Table S1.** Promoter DNA Constructs.

| Promoter | Template | Upstream primer | Downstream primer | Duplex (bp) and position relative to start site (+1)<sup>a</sup> | Experiment used in |
| --- | --- | --- | --- | --- | --- |
| $\lambda P_R$ (-59 to +34)<sup>b</sup> | Unlabeled $\lambda P_R$ | Up | Down | 192 (-128 to +64) | Permanganate footprinting kinetics |
| Cy3(+14) $\lambda P_R$ <sup>c</sup> | Unlabeled $\lambda P_R$ | Up | Cy3(+14) Down | 142 (-128 to +14) | Fluorescence kinetics |
| Cy5(+14) $\lambda P_R$ <sup>c</sup> | Unlabeled $\lambda P_R$ | Up | Cy5(+14) Down | 142 (-128 to +14) | |
| Cy3(-100) $\lambda P_R$ <sup>c</sup> | Unlabeled $\lambda P_R$ | Cy3(-100) Up | Down | 164 (-100 to +64) | |
| Cy5(-100) $\lambda P_R$ <sup>c</sup> | Unlabeled $\lambda P_R$ | Cy5(-100) Up | Down | 164 (-100 to +64) | |
| Cy3(+14)-Cy5 (-100) $\lambda P_R$ <sup>c</sup> | Unlabeled $\lambda P_R$ | Cy5(-100) Up | Cy3(+14) Down | 114 (-100 to +14) | |
| Cy5 (+14)- Cy3 (-100) $\lambda P_R$ <sup>c</sup> | Unlabeled $\lambda P_R$ | Cy3(-100) Up | Cy5(+14) Down | 114 (-100 to +14) | |
| <sup>a</sup> Dye-labeled DNAs have 12 base (upstream) and/or 8 base (downstream) ssDNA termini |  |  |  |  |  |
| <sup>b</sup> SI Ref. 13 <sup>c</sup> SI Ref. 1 |  |  |  |  |  |

**Table S2.**
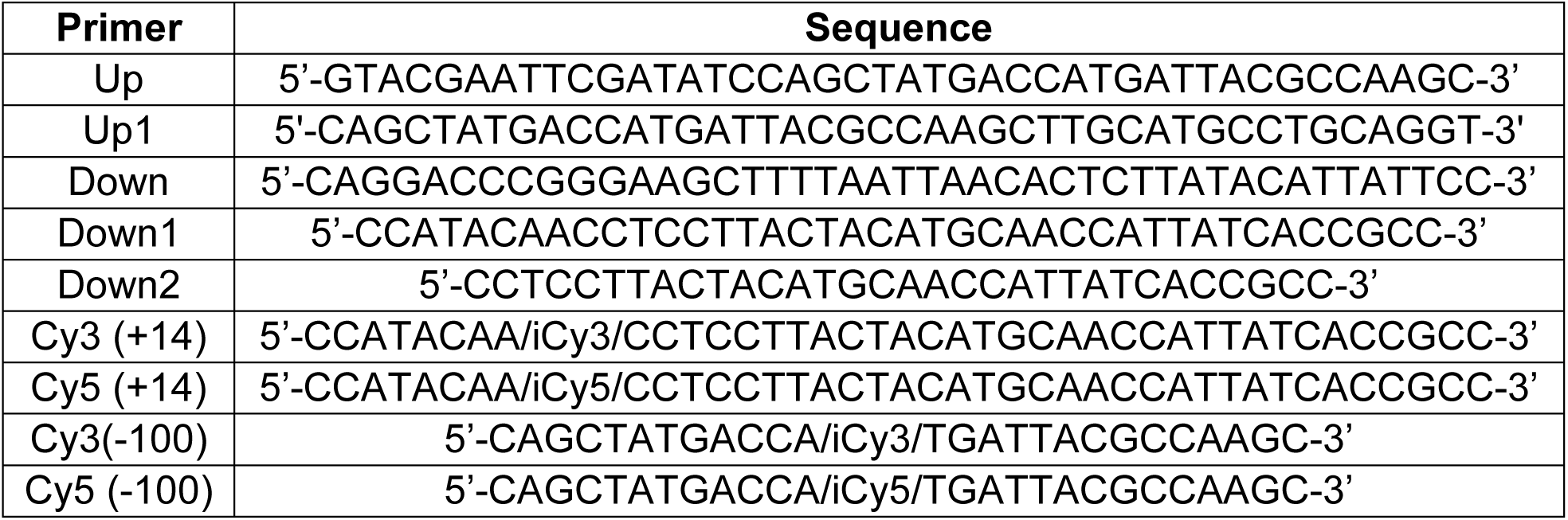
PCR Primer Sequences.

**Table S3.** Comparison of Observed and Predicted *τ*_*i,obs*_ Values

|  | **Observed $1/\tau_{i,obs}$ from**<br>**FRET, PIFE Fits ($\pm 1$ SD)** | **Predicted $1/\tau_{i,pred}$ from**<br>**Rate Constants (Table 3)<sup>a</sup>** |
| --- | --- | --- |
| $1/\tau_{1,obs}$ | $16 \pm 12 \text{ s}^{-1}$ | $10.5 \pm 0.6 \text{ s}^{-1}$ |
| $1/\tau_{2,obs}$ | $2.6 \pm 1.5 \text{ s}^{-1}$ | $2.4 \pm 0.1 \text{ s}^{-1}$ |
| $1/\tau_{3,obs}$ | $0.21 \pm 0.18 \text{ s}^{-1}$ | $0.12 \pm 0.01 \text{ s}^{-1}$ |
| $1/\tau_{4,obs}$ | $0.020 \pm 0.019 \text{ s}^{-1}$ | $0.010 \pm 0.002 \text{ s}^{-1}$ |

<sup>a</sup> Predicted using Eqs. S3-S6 and the finding from the simulation in Fig. 7 that in the time range of most interest (1 s -10 s), $C_{eq} = ([P]_{eq} + [R]_{eq}) \approx [P]_{tot}$ , the total promoter concentration.

**Table S4.** Rate Constants, Amplitudes of FRET and PIFE Kinetics Experiments (Fig. 6)

|  | FRET |  |  |  | PIFE |  |  |  |
| --- | --- | --- | --- | --- | --- | --- | --- | --- |
|  | Cy5 +14 |  | Cy5 -100 |  | Cy3 +14 | Cy5 +14 | Cy3 -100 | Cy5 -100 |
|  | Initial<sup>a</sup> | Refined<sup>b</sup> | Initial<sup>a</sup> | Refined<sup>b</sup> | Refined<sup>b</sup> | Refined<sup>b</sup> | Refined<sup>b</sup> | Refined<sup>b</sup> |
| **k<sub>1</sub>(M<sup>-1</sup> s<sup>-1</sup>)** | **2.8 x10<sup>8</sup>** | **3.0 x10<sup>8</sup>** | **3.1 x10<sup>8</sup>** | **2.8 x10<sup>8</sup>** | **3.4 x10<sup>8</sup>** | **3.0 x10<sup>8</sup>** | **3.4 x10<sup>8</sup>** | **2.8 x10<sup>8</sup>** |
| **k<sub>-1</sub> ( s<sup>-1</sup>)** | **40** | **48** | **44** | **43** | **42** | **43** | **40** | **44** |
| **k<sub>2</sub> ( s<sup>-1</sup>)** | **13** | **14** | **15** | **13** | **14** | **14** | **12** | **14** |
| **k<sub>-2</sub> ( s<sup>-1</sup>)** | **6.7** | **7.5** | **7.4** | **7.1** | **7.1** | **7.1** | **7.0** | **6.6** |
| **k<sub>3</sub> ( s<sup>-1</sup>)** | **2.2** | **2.3** | **2.5** | **2.4** | **2.5** | **2.6** | **2.4** | **2.5** |
| **k<sub>-3</sub> ( s<sup>-1</sup>)** | **1.7** | **1.9** | **1.9** | **1.7** | **1.6** | **1.9** | **1.6** | **1.6** |
| **k<sub>4</sub> ( s<sup>-1</sup>)** | **0.10** | **0.098** | **0.088** | **0.096** | **0.096** | **0.090** | **0.11** | **0.097** |
| **k<sub>-4</sub> ( s<sup>-1</sup>)** | **0.10** | **0.11** | **0.088** | **0.091** | **0.098** | **0.10** | **0.092** | **0.098** |
| **k<sub>5</sub> ( s<sup>-1</sup>)** | **0.040** | **0.040** | **0.036** | **0.038** | **0.041** | **0.039** | **0.042** | **0.044** |
| **A<sub>I1E</sub>** | **0.36** | **0.37** | **0.61** | **0.68** | **2.4** | **2.9** | **2.1** | **1.5** |
| **A<sub>I1M</sub>** | **0.25** | **0.29** | **0.20** | **0.20** | **2.0** | **1.7** | **2.5** | **3.0** |
| **A<sub>I1L</sub>** | **0.51** | **0.69** | **1.2** | **1.1** | **1.5** | **2.1** | **0.83** | **0.33** |
| **A<sub>OC</sub>** | **1.2** | **1.3** | **1.2** | **1.2** | **1.1** | **1.2** | **0.67** | **0.79** |
| **RMS <sup>c</sup>** | **0.014** | **0.0081** | **0.01** | **0.009** | **0.014** | **0.023** | **0.019** | **0.054** |

<sup>a</sup> Initial Kintek fit with no drift correction
<sup>b</sup> Refined Berkley Madonna fit using Eq. S7 with a linear drift correction as in Eq. S2 and Kintek average rate constants $\pm 10\%$ .
<sup>c</sup> RMS (root mean square) error calculated by Berkeley-Madonna for fits in Fig 6.

**Table S5.**
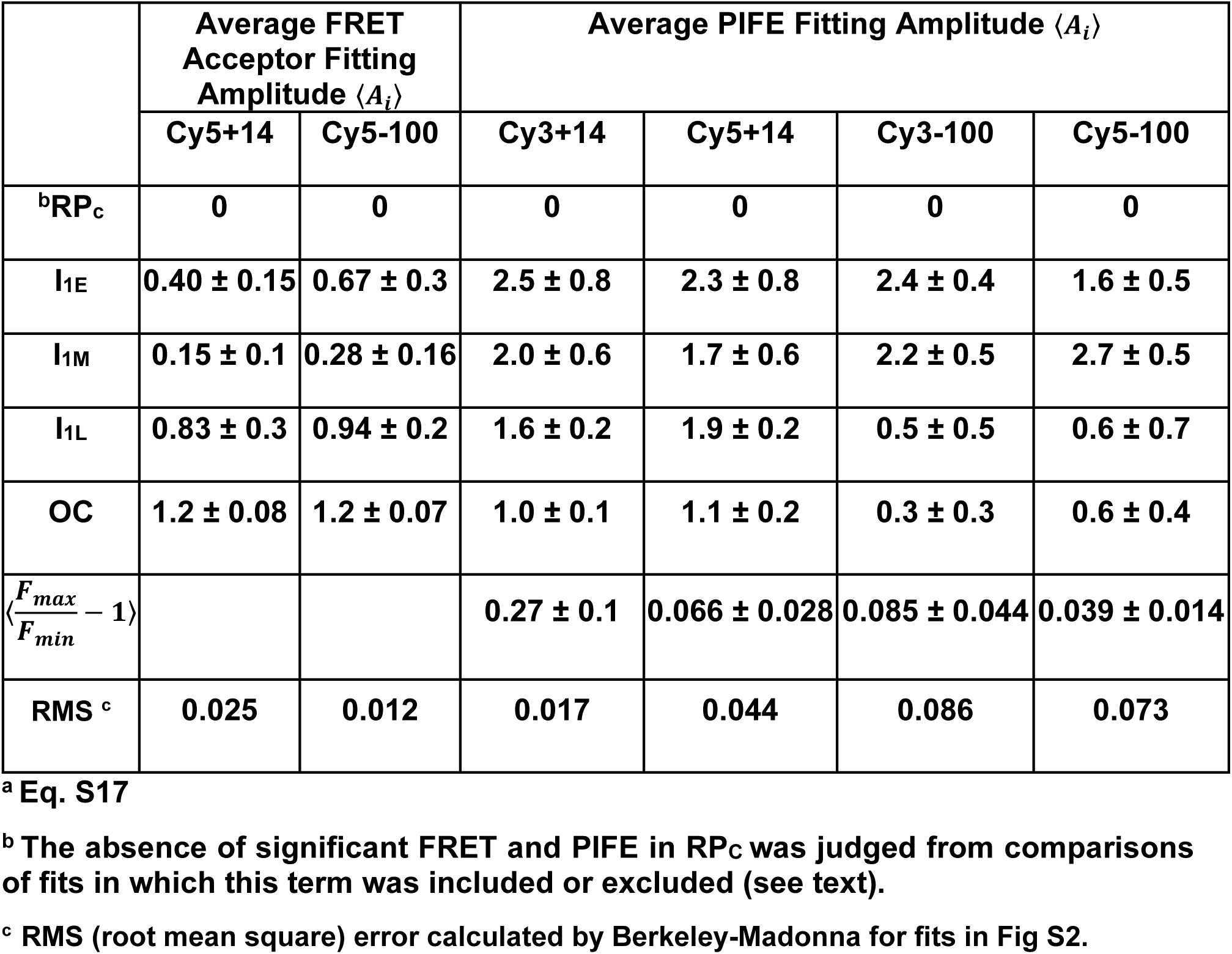
Average FRET Acceptor and PIFE Fitting Amplitudes ⟨*A*_*i*_⟩ of CC Intermediates and OC and Conversion Factors 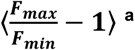

